# Ancestral reconstruction of the MotA stator subunit reveals that conserved residues far from the pore are required to drive flagellar motility

**DOI:** 10.1101/2022.10.17.512626

**Authors:** Md Imtiazul Islam, Pietro Ridone, Angela Lin, Katharine A Michie, Nicholas J Matzke, Georg Hochberg, Matthew AB Baker

## Abstract

The bacterial flagellar motor (BFM) is a rotary nanomachine powered by the translocation of ions across the inner membrane through the stator complex. The stator complex consists of two membrane proteins: MotA and MotB (in H^+^ powered motors), or PomA and PomB (in Na^+^ powered motors). In this study we used ancestral sequence reconstruction (ASR) to probe which residues of MotA correlate with function and may have been conserved to preserve motor function. We reconstructed ten ancestral sequences of MotA and found four of them were motile in combination with contemporary *E. coli* MotB and in combination with our previously published functional ancestral MotBs. Sequence comparison between wild-type (WT) *E. coli* MotA and MotA-ASRs revealed 30 critical residues across multiple domains of MotA that were conserved among all motile stator units. These conserved residues included pore-facing, cytoplasm-facing and MotA-MotA intermolecular facing sites. Overall, this work demonstrates the role of ASR in assessing conserved variable residues in a subunit of a molecular complex.

## Introduction

The bacterial flagellar motor (BFM) is a rotary molecular motor responsible for motility in many bacteria aiding their survival and pathogenicity. The BFM is one of the largest (11 MDa), dynamically self-assembled, and membrane-spanning molecular machines in bacteria, requiring more than a dozen different proteins to assemble and function (Sowa and Berry, 2008; Beeby et al., 2020). It contains a cytoplasmic rotor and a ring of stator units that surround the rotor (Berg, 2003; Nakamura and Minamino, 2019). The stator powers the motor by transporting ions across the cell membrane, generating torque in interaction with the rotor, which is coupled to the flagellar filament (Minamino and Imada, 2015).

The stator of the BFM is a complex of two membrane proteins: MotA and MotB in H^+^ powered motors in *Escherichia coli*, and PomA and PomB in Na^+^ powered motors in *Vibrio alginolyticus* (Yorimitsu and Homma, 2001; Berg, 2003). MotA and MotB form a heterodimer with a stoichiometry of MotA_5_MotB_2_ that creates an ion channel at the rotor interface (Deme et al., 2020; Santiveri et al., 2020). The MotA subunit contains four transmembrane (TM) helices (TMH1–TMH4), two short periplasmic loops, and a cytoplasmic loop between TMH2 and TMH3 and the C-terminal cytoplasmic tail (Zhou and Blair, 1997; Hu et al., 2022). The cytoplasmic loop contains highly conserved charged residues that interact with the rotor protein FliG (Zhou et al., 1998a). Conversely, the MotB subunit contains a single TM helix that lines the ion pathway alongside the TMH3 and TMH4 helices of MotA (Kojima and Blair, 2001). Adjacent to the pore-lining MotB TM domain is a periplasmic region containing a plug that can block the pore with a spanner-like mechanism (Homma et al., 2021). A peptidoglycan-binding domain (PGD) anchors the stator through binding to the PG-layer (Hosking et al., 2006). Interactions between MotA and the rotor trigger a structural change of the PGD and the plug region, that allows binding of PGD to the peptidoglycan (PG) layer, unplugging of the pore and activating the stator complex to allow ion flow and torque generation (Kojima et al., 2018).

The TM domains of both MotA and MotB contain a series of conserved residues that are known to be important for flagellar function, ion selection and transfer (Asai et al., 2000a; Ito et al., 2005). The aspartic acid residue 32 at the TM region of MotB (MotB-D32) acts as a universally conserved ion-binding site (Zhou et al., 1998b; Che et al., 2008), and the D32s from both MotB subunits are exposed to the pentameric MotA ring at the subunit interface in proximity of MotA-T209 (Deme et al., 2020). Among other reported conserved and critical residues of the stator, MotA-R90 and MotA-E98 contribute to motor rotation with rotor protein FliG (Zhou and Blair, 1997; Takekawa et al., 2014), MotA-P173 and MotA-P222 interact with adjacent MotA monomers stabilizing the structure (Braun et al., 1999; Deme et al., 2020), and MotA-A180 is involved in ion specificity with MotB-Y30 (Biquet-Bisquert et al., 2021). Over the years, many mutagenesis techniques have been used to study the structural residues of BFM, such as random mutagenesis (Blair and Berg, 1990), tryptophan-scanning mutagenesis (Sharp et al., 1995), cysteine-scanning mutagenesis (Asai et al., 2000b), site-directed mutagenesis (Nakamura et al., 2009), photo cross-linking and disulfide cross-linking (Terashima et al., 2021). In this study, we used ancestral sequence reconstruction (Hochberg and Thornton, 2017; Garcia and Kaçar, 2019; Dube et al., 2022) to generate hypothetical ancestral states from phylogenies of extant proteins. We reconstructed and synthesized ten ancestral MotAs to determine which residues of MotA were conserved throughout the evolution of BFM and were essential for BFM function.

## Results

### Phylogeny of MotA and the selected Nodes for resurrection

Two phylogenetic trees were calculated for two separate multiple sequence alignments comprising of 178 and 264 MotA homologous sequences (Fig. 1) Ten nodes across both phylogenies were selected for ancestral sequence reconstruction (ASR), focusing on a mixture of older and younger ancestors across known sodium- and proton-motile clades. Among the ten selected nodes, ASR180, ASR220, ASRN41, and ASR333 had descendants that only belonged to proteobacteria, whereas ASR244, ASRN65, ASR259, ASR266, ASR332, and ASR440 had descendants that also included terrabacteria (actinobacteria, firmicutes), aquificae, and spirochaetes. Pairwise internode distance and pairwise percent coidentity between each node is shown in Supplementary Fig. 1 & 2; posterior probability across each site for ASRs is shown in Supplementary Fig. 3. We performed structural prediction for all reconstructed MotA-ASRs using ColabFold (Mirdita et al., 2021, 2022) prior to synthesis to compare predicted structures of the ASRs with existing known MotA structures (PDBIDSupplementary Fig. 4-6).

**Figure 1:**
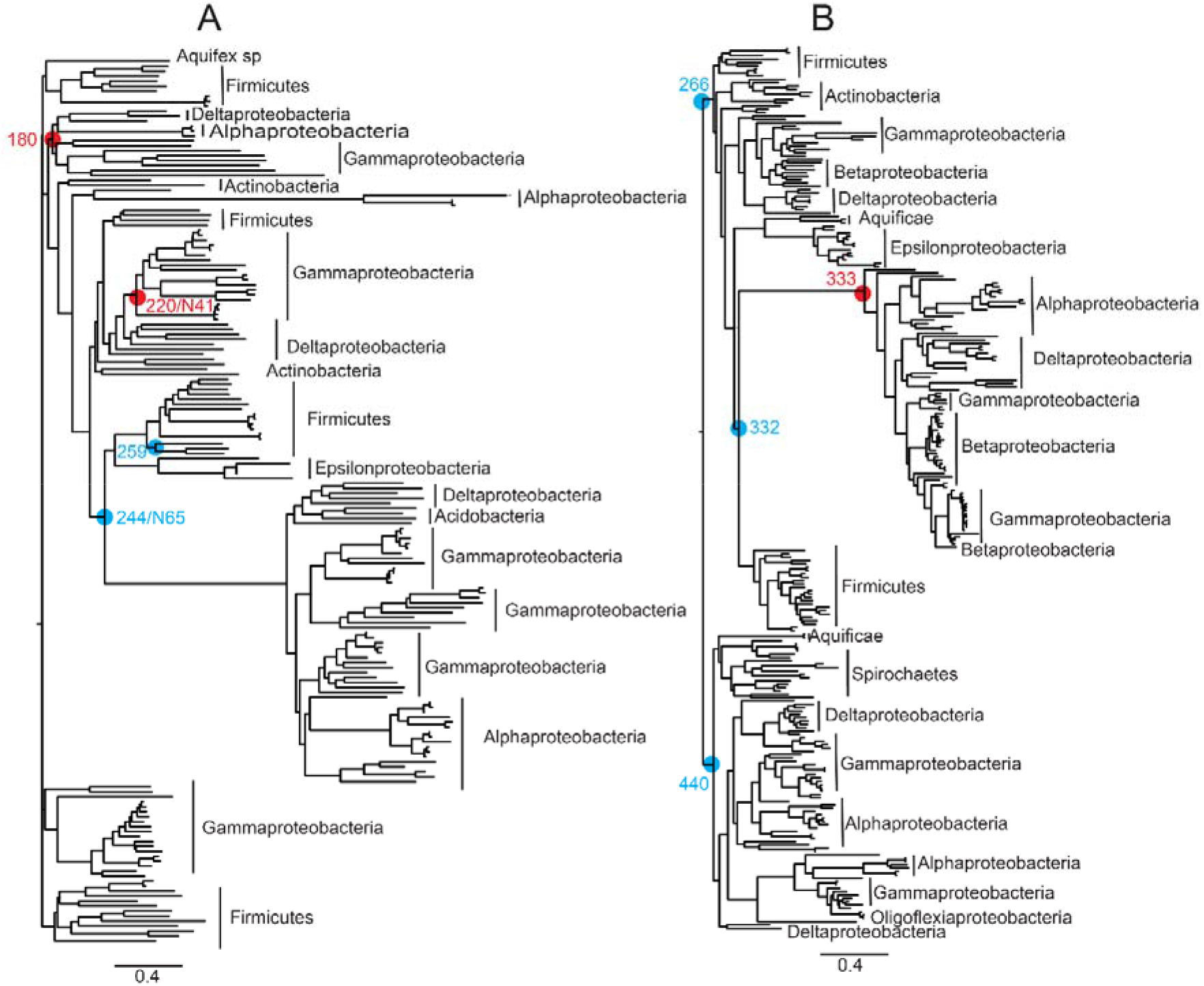
Phylogenetic trees of 178 and 264 *motA* homologs. Ten nodes across both phylogenies were selected for synthesis (red circles: motile MotA-ASRs; blue circles: non-motile MotA-ASRs). (A) Neighbour-joining phylogeny generated using QuickTree (Howe et al., 2002) for 178 *motA* sequences. (B) Maximum-likelihood tree generated using RAxML (Stamatakis, 2014) for 264 *motA* sequences. Scale bar indicates substitutions per site. Newick files for the phylogenies, sequence alignments, and ASR quality measurements are available in the Supplementary Material.

### Functional characterisation of MotA-ASRs

We evaluated the functional properties of all ten reconstructed MotA-ASR sequences by testing their motility through plasmid-based expression in a Δ*motA* strain. Overall, we characterised semi-solid agar swimming, cell free swimming, and tethered-cell rotation in different concentrations of sodium. Swim plate assay results indicated that three out of the ten MotA-ASRs (ASR180, ASR220, and ASR333) were able to restore motility in combination with contemporary WT *E. coli* MotB in Δ*motA* RP437 (Fig. 2A). ASR180, ASR220 and ASR333 were also motile in the free-swimming assay with mean speeds of 1.6 ± 0.8, 3.0 ± 1.3, and 2.7 ± 1.1 μm/s, respectively (Fig. 2B). In comparison, the Na^+^ (*pomApotB*) and H^+^ (*motAmotB*) powered swimming controls had an average speed of 3.3 ± 1.3, and 5.2 ± 1.5 μm/s respectively. Both motile and non-motile MotA-ASRs exhibited similar growth rates in combination with WT *E. coli* MotB in Δ*motA* RP437 (Supplementary Fig. 7).

**Figure 2:**
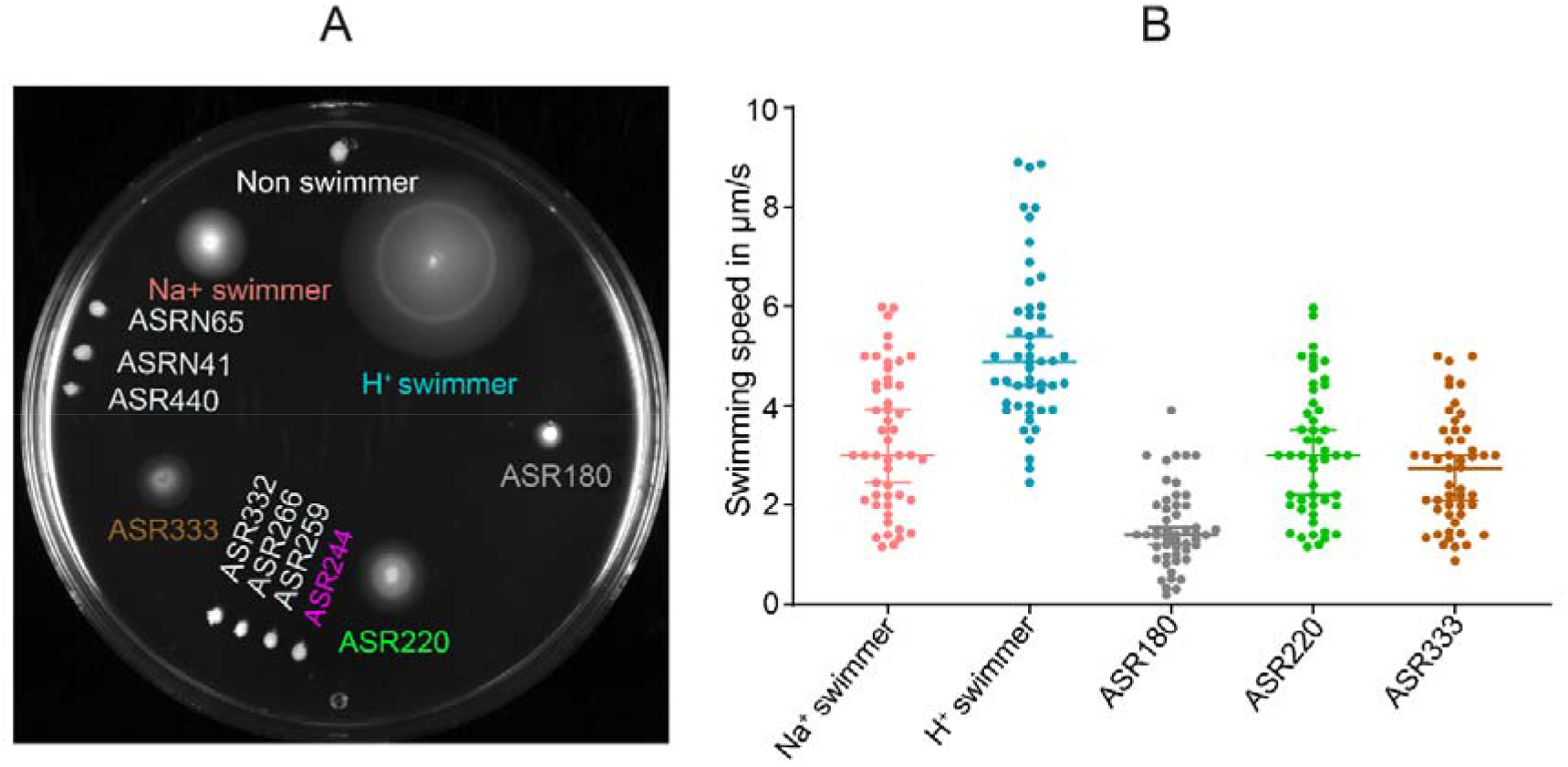
Functional properties of MotA-ASRs. (A) Swim plate assay (LB + 0.25% agar) of Mot-ASRs, H+ powered (*motAmotB*) and Na+ powered (*pomApotB*) control swimmers and a control non-swimmer (empty vector, pBAD33) on LB swim agar. After 14 h incubation at 30°C, ASR180, ASR220 and ASR333 produced swimming halos (coloured labels) while the other 7 MotA-ASRs did not produce swimming halos (white labels) (B) Swimming speeds of three functional MotA-ASRs (ASR180, ASR220 and ASR333 and the control H^+^/Na^+^ swimmers in a free-swimming assay. Colours in B match labels in A. The speed of individual cells was measured in μm/s and is shown with the scattered plot; median speeds are indicated by the horizontal line with a 95% confidence interval (n = 50 cells per ASR).

### Ionic power source driving rotation in functional MotA-ASRs

We evaluated the ionic power source (H^+^ or Na^+^) of our motile MotA-ASRs in low Na^+^ conditions (~1 mM) using a minimal swim plate without tryptone or yeast extract and where NaCl had been substituted with KCl (Islam et al., 2020; Ridone et al., 2022). Here, we checked whether the tested strains could generate swim rings or not in the low sodium environment. The result showed that the H^+^-powered control (*motAmotB*) and each of the three motile MotA-ASRs (ASR180, ASR220 and ASR333) produced swimming rings on the minimal swim plate (Fig. 3A). In contrast, the Na^+^ powered control (*pomApotB*) and other non-motile MotA-ASRs did not produce any swimming rings (Fig. 3A). Tethered cell assay results agreed: the rotational speeds of ASR180, ASR220, ASR333 and the H^+^ powered control remained constant as sodium concentration was varied and were 2.0 ± 0.9 Hz, 2.9 ± 0.7 Hz, 2.75 ± 0.7 Hz and 3.8 ± 0.7 Hz respectively (Figure 3B). However, the rotation speed of the Na^+^ powered control increased from stationary (0 Hz) to 2.8 ± 0.7 Hz with the increase of external sodium concentration from 0 to 85 mM (Fig. 3B). H^+^ conductivity was further evaluated by examining growth curves for all MotA-ASRs when combined with plug-deleted MotB_Δ521-70_ Growth of WT *E. coli* MotA and the three motile MotA-ASRs (ASR180, ASR220 and ASR333) was inhibited when combined with MotB_Δ51-70_ (Figure 3CD and Supplementary Fig. 8). In contrast, growth was not inhibited when non-motile MotA-ASRs were combined with plug-deleted MotB_Δ51-70_ (Figure 3D and Supplementary Fig. 8).

**Figure 3:**
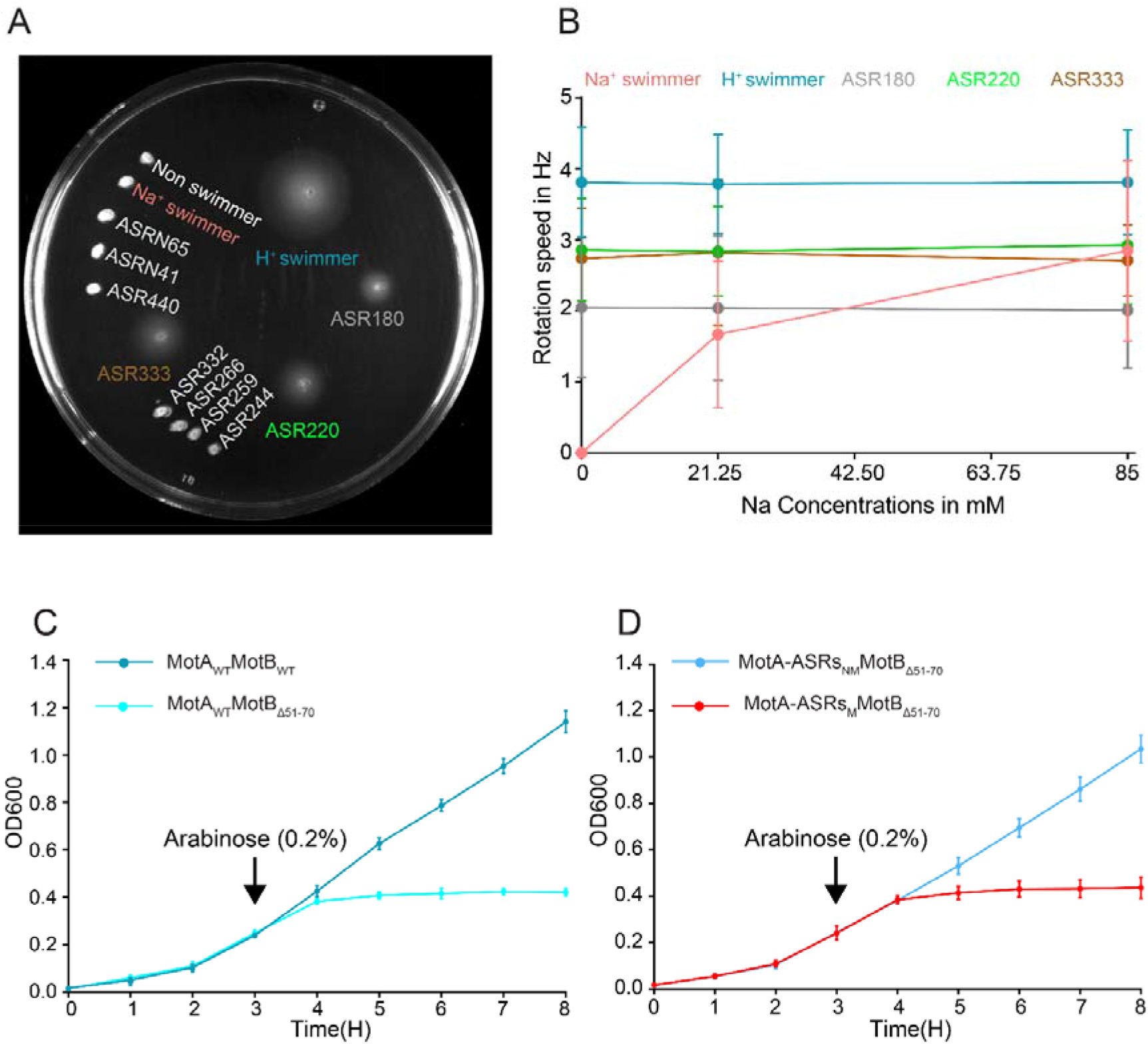
Characterization of the ion selectivity of MotA-ASRs. (A) Minimal swim plate assay with KCl instead of NaCl (~1 mM [Na^+^]) for all MotA-ASRs and the Na^+^ and H^+^ powered control. After 18 hours incubation at 30°C, control Na^+^ swimmer (*pomApotB*) and all non-functional MotA-ASRs were non-motile. In contrast, control H^+^ swimmer (*momAmotB*) and the tested functional MotA-ASRs (ASR180, ASR229, ASR333) were motile and produced swimming rings. (B) The rotational speed of MotA-ASRs and the Na^+^ and H^+^ powered controls were calculated in the presence of 0, 21.25, and 85 mM NaCl from the tethered cell assay (measured in Hz). The average rotational speed of all tested strains (mean ± SD, *n* = 50 cells per ASR) is shown in the line graph. Colours in B match labels in A. (C) Growth curves of wild type MotA (MotA_WT_) in combination with wild type MotB (MotB_WT_) and plug deleted MotB (MotB_Δ51-70_). (D) Growth curves of motile MotA-ASRs (MotA-ASRsM) and non-motile MotA-ASRs (MotA-ASRsNM) in combination with plug deleted MotB (MotB_Δ51-70_). The averaged growths (mean ± SD) of three motile (red line) and six non-motile MotA-ASRs (blue line) were showed. Full data for each ASR is shown in Supplementary Figure 8.

### Functional compatibility of MotA-ASRs with MotB-ASRs and *Aquifex* MotBs

We tested multiple stator combinations by examining a 150 element swim-plate array. We examined all combinations of 10 MotA-ASRs with 15 ancestral MotBs consisting of our previously published 13 MotB-ASRs and 2 *Aquifex* MotBs (Islam et al., 2020) in a Δ*motAmotB* strain to evaluate their intercompatibility. Four out of ten of the MotA-ASRs (ASR180, ASR220, ASR333 and ASRN41) were compatible with several MotB-ASRs and formed functional stator units. The expression of MotA-ASR220 restored motility with each of the 13 MotB-ASRs. In contrast, MotA-ASR180, MotA-ASR333, and MotA-N41 were able to restore motility with 11, 11, and 1 of 13 MotB-ASRs, respectively (Fig. 4). The other six MotA-ASRs, ASR244, ASR259, ASR266, ASR332, ASR440, and ASRN65 did not show compatibility with any of the 13 MotB-ASRs and were non-motile (Fig. 4). None of the ten MotA-ASRs restored motility when combined with either of the two Aquifex MotBs (Fig. 4).

**Figure 4:**
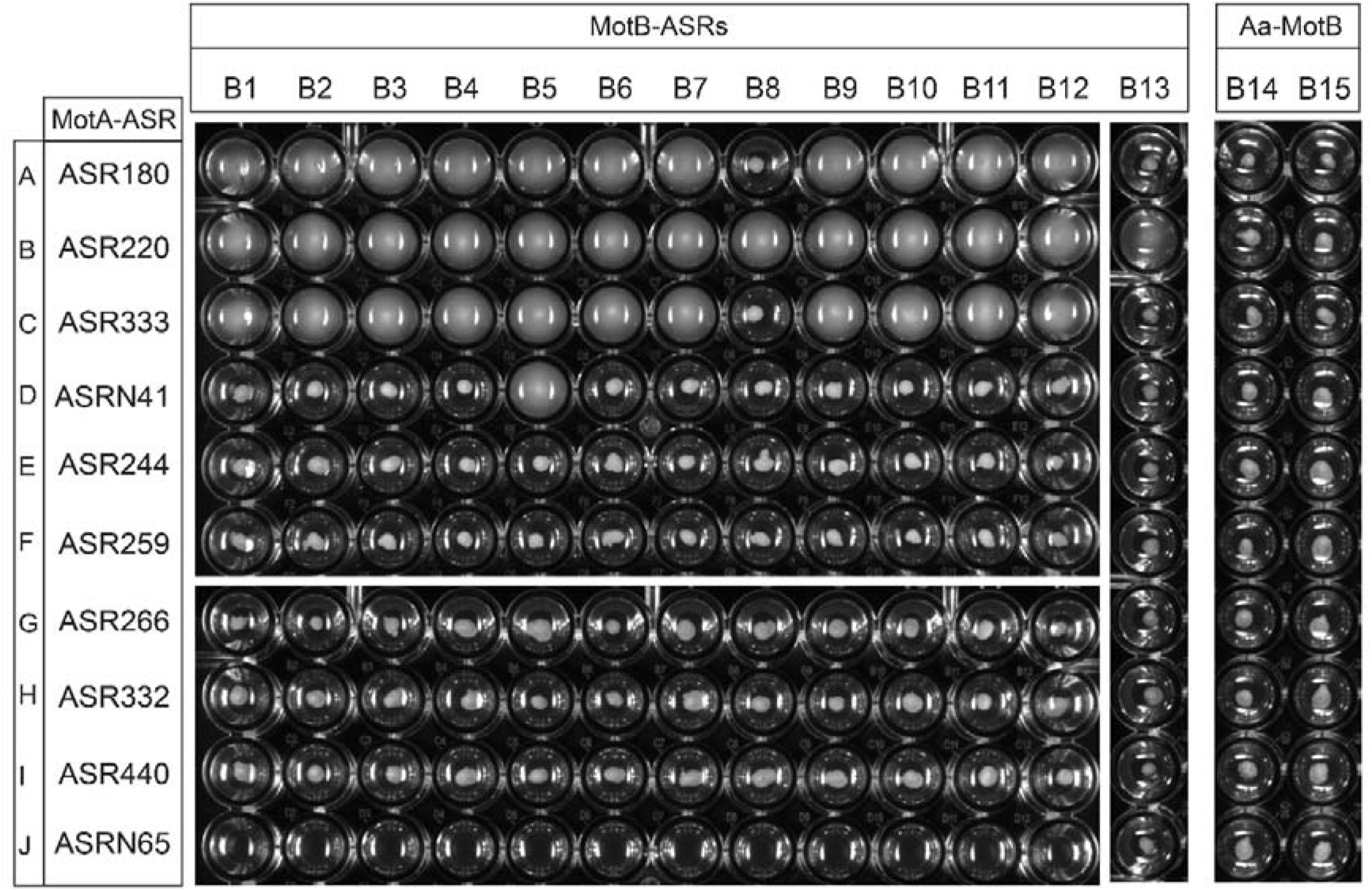
Compatibility between MotA-ASRs and MotB-ASRs. Swim plate assay in 96 well plate showing functional compatibility of MotA-ASRs with MotB-ASRs from (Islam et al., 2020). Each well contains 0.25% LB swim agar and the plate was incubated 14 hours at 30°C after inoculation of the test combination. A-J rows contain MotA-ASRs, and B1-B13 and B14-B15 columns contain tested MotB-ASRs and *Aquifex aeolicus* (Aa) MotBs, respectively. Row-A: MotA180; Row-B: MotA220; Row-C: MotA333; Row-D: MotAN41; Row-E: MotA244; Row-F: MotA259; Row-G: MotA266; Row-H: MotA332; Row-I: MotA440; and Row-J: MotAN65; Col-B1: MotB758; Col-B2: MotB759; Col-B3: MotB760; Col-B4: MotB765; Col-B5: MotB908; Col-B6: MotB981; Col-B7: MotB1024; Col-B8: MotB1170; Col-B9: MotB1239; Col-B10: MotB1246; Col-B11: MotB1457; Col-B12: MotB1459; Col-B13: MotB1501; Col-B14: Aa MotB1 and Col-B15: Aa MotB2.

### Conservation of residues between WT *E. coli* - MotA and the motile MotA-ASRs

We examined residue conservation between WT *E. coli* MotA and four motile MotA-ASRs (ASR180, ASR220, ASR333 and ASRN41) by performing a multiple sequence alignment. The alignment of WT *E. coli* MotA and four motile MotA-ASRs (Supplementary Fig. 9) showed that 81 of the 295 sites (27.5%) in all motile MotA-ASRs were identical, 96 sites (32.5%) were similar (i.e., were replaced, relative to WT, with a residue with similar biochemical properties), and 62 sites (21%) were a mixture of both similar and distinct sites across all the motile MotA-ASRs when compared with *E. coli* MotA. The remaining 56 (19%) of 295 residues were distinct when compared with *E. coli*-MotA at the same position (i.e., were substitutions of amino acids with notable changes in hydrophobicity, polarity, or size, in all four functional ASRs) (Fig. 5). Sequence comparison and conservation was mapped onto an Alphafold model (Jumper et al., 2021) of the WT *E. coli* MotA_5_MotB_2_ complex (Fig. 5). The pLDDT and PAE plots of the *E. coli* MotAB Alphafold model, with colour-coded confidence overlay onto the structure are provided in Supplementary Fig. 10. The RMSDs for each chain of the *E. coli* MotAB Alphafold model in comparison with each of 6YKP, 6YKM and 6YSL (reported MotAB structures of *C. jejuni* and *B. subtilis*) are provided in Supplementary Table 1.

**Figure 5:**
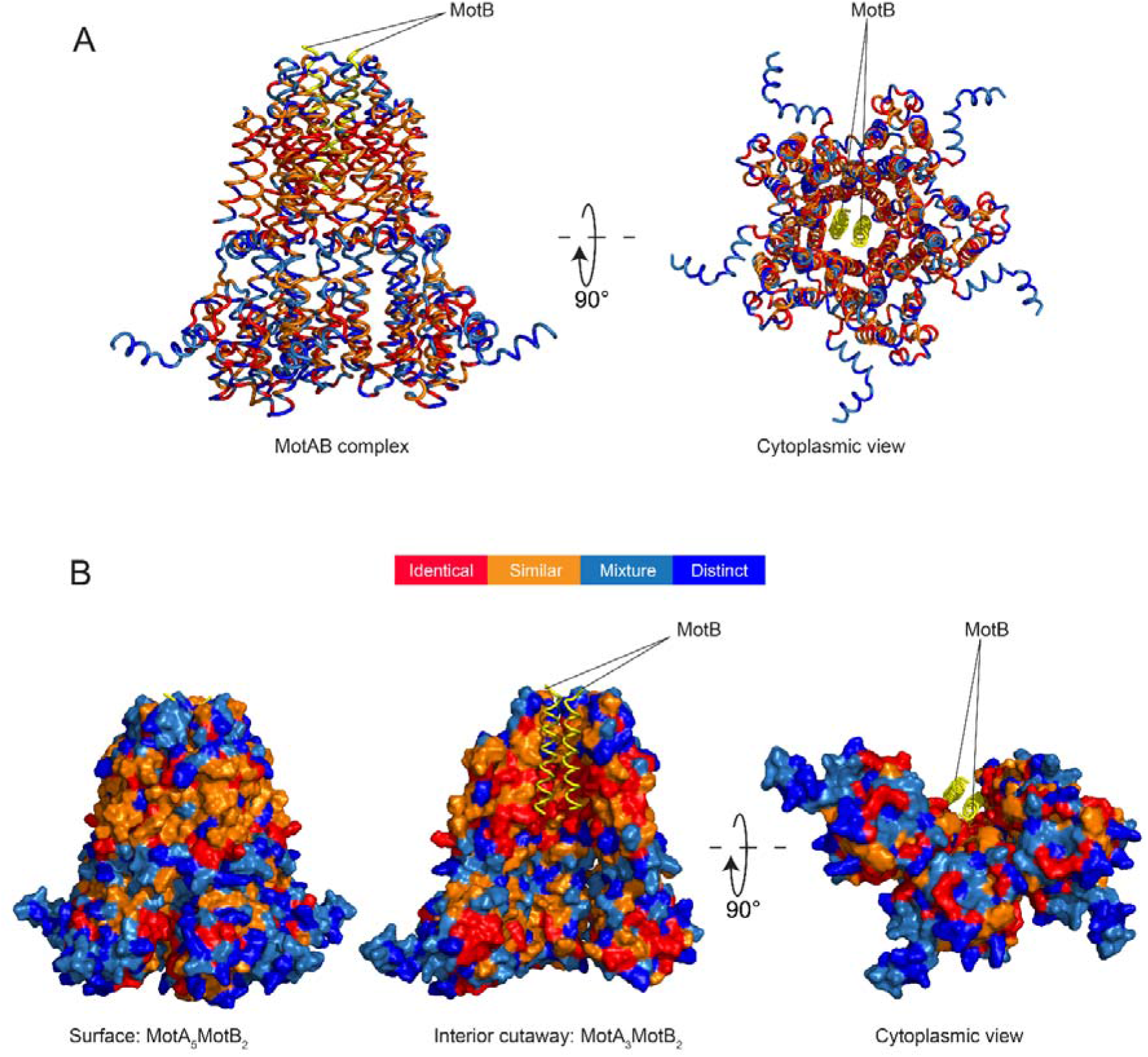
Residue conservation and variation between WT and reconstructed MotAs mapped onto an Alphafold model of *E. coli* MotA_5_MotB_2_. (A) Cartoon model shows the identical residues across the functional MotA-ASRs (red), similar (orange), mixed (light blue) and distinct (dark blue), visualised from the side (left) and from the cytoplasm (right). MotB is coloured yellow. (B) Surface model of MotAB complex (A5B2) with MotB is shown as cartoon. Side view showing the external surface conservation of MotA (Left), interior cutaway of the MotA_3_B_2_ complex showing the internal conservation (middle), and bottom cytoplasmic view of the interior cutaway of the MotA_3_B_2_ complex (right).

### Determination of the residues critical for function from comparison of wild-type MotA with both motile and non-motile MotA-ASRs

Sequence comparison between our resurrected MotA-ASRs and WT *E. coli* MotA revealed several residues critical to MotA function. Sequence comparison highlighted that 30 (10%) of the 295 residues in the motile MotA-ASRs differed from the corresponding residues in the non-motile MotA-ASRs, but were identical to the corresponding residues of WT *E. coli* MotA (Supplementary Fig. 11). We located each of the 30 identified critical residues on our *E. coli* MotA_5_MotB_2_ structural model and observed that these critical residues were distributed throughout the MotA domains (Fig. 6). Among the 30 residues, four were found in transmembrane helix 2 (TMH2), eight in cytoplasmic helix 1-3 (CPH1-3), four in transmembrane helix 3 (TMH3), six in transmembrane helix 4 (TMH4) and eight in cytoplasmic helix 4 (CPH4) (Fig. 6 and Supplementary Fig. 11). We measured the intermolecular distances between our conserved residues and several previously reported residues that are essential to BFM function (Supplementary Table 2) and observed that these residues were in close proximity (ranging from 2.3-7.1 Å). In particular, we observed the proximity of MotA-F45 with MotA-I54 and V168, MotA-D128 with MotA-E262, MotA-Y217 with MotA-P173 and MotB-W26, and MotA-L223 with MotA-L64 (Supplementary Fig. 12).

**Figure 6:**
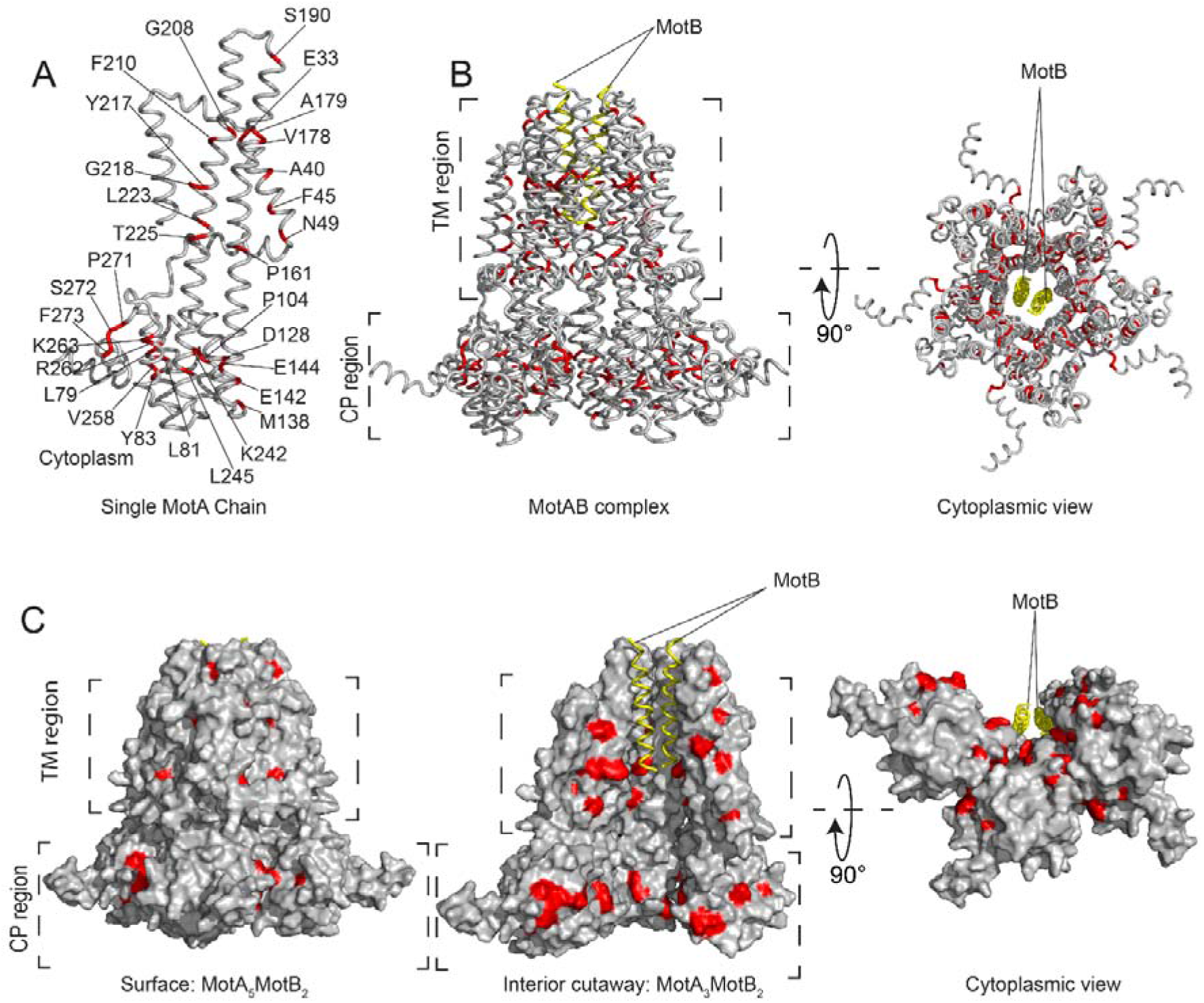
Revealed critical residues of MotA mapped onto an Alphafold model of *E. coli.*. (A) 30 critical residues from functional and nonfunctional MotA-ASRs are highlighted in red (conserved) with their identity on an isolated MotA chain. (B) The distribution and location of the 30 critical residues are showed on the complete MotA_5_MotB_2_ (side view and bottom cytoplasmic view). (C) Location of 30 residues labelled in (A) coloured in red on the surface and interior of the MotA_5_MotB_2_ complex (left) side view of the outside of the surface with critical residues, (centre), interior view with MotA trimer and MotB dimer and bottom cytoplasmic view (right).

### Experimental verification of the role of the selected conserved residues in flagellar function

We then tested whether the introduction of point mutations at specific residues could rescue motility in non-motile MotA-ASRs, or disrupt motility in either motile MotA-ASRs or WT *E. coli* MotA. We selected six MotA residues (A40, V178, A179, Y217, G218 and R262) of interest based on previous reports (Garza et al., 1996; Deme et al., 2020; Biquet-Bisquert et al., 2021). Replacement of residues at selected positions in WT *E. coli* and motile MotA-ASRs with residues from non-motile MotA-ASRs at the same site showed that V178I and A179G point mutations did not disrupt motility, but the Y217N mutation completely disrupted motility (Figure 7A and Supplementary Table 3). In contrast, the reintroduction of residues from motile MotA-ASRs at the selected position of non-motile MotA-ASRs showed that none of the tested mutations (G40A, I178V, G179A, N217Y, L/M/V/A218G, and K262R), individually or in combination, were able to rescue motility (Figure 7B and Supplementary Table 3).

**Figure 7:**
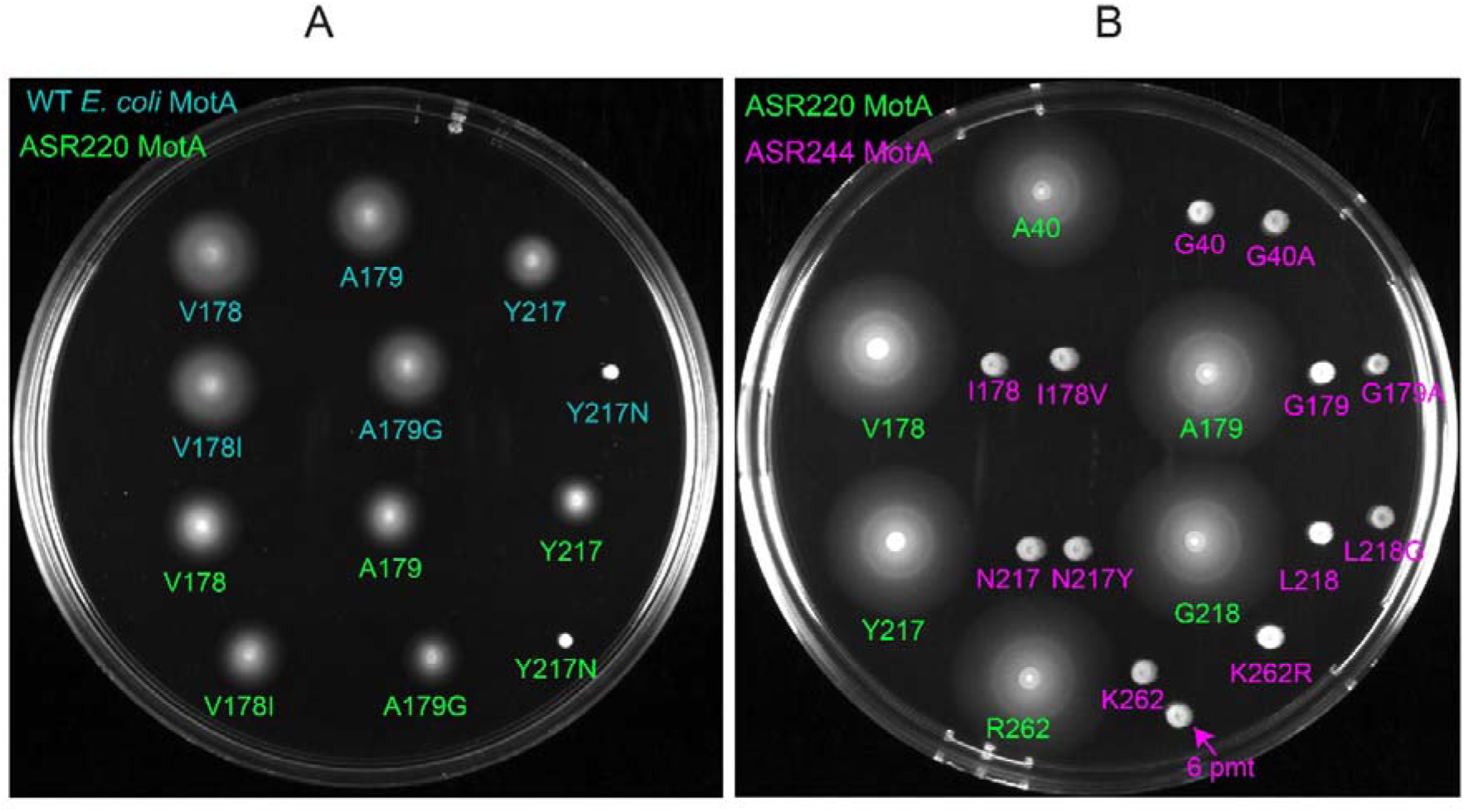
Role of the point mutations in disrupting or rescuing motility of MotA. (A) Swim plate assay (LB + 0.25% agar) comparing motility of WT *E. coli*-MotA (coloured in deep teal) and the functional MotA-ASR220 (coloured in green): V178, A179 and Y217 with their point mutants: V178I, A179G, and Y217N. (B) Swim plate assay (LB + 0.25% agar) comparing motility of functional MotA-ASR220 (coloured in green): A40, V178, A179, Y217, G218 and R262, non-functional MotA-ASR244 (coloured in magenta): G40, I178, G179, N217, L218 and K262 and point mutants of non-functional MotA-ASR244: G40A, I178V, G179A, N217Y, L218G and K262R. Magenta arrow indicates strain with all 6 point mutations (6 pmt).

## Discussion

This study 1) verifies the roles of MotA residues previously reported as crucial, 2) provides evidence as to which regions of MotA are variable, and 3) reveals new residues that are conserved and essential for flagellar function. First, several of our identified conserved residues, including MotA-V178, MotA-A178, and MotA-Y217 (Supplementary Fig. 7), are located near the pore in transmembrane helices 3 and 4 (TM3/TM4) and these have been widely reported to be important for stator function (Zhou and Blair, 1997; Braun et al., 1999; Kim et al., 2008; Nakamura et al., 2009; Deme et al., 2020). Other conserved residues are found in the cytoplasmic region and in TM helix-2 (TM2), and a number of those, such as MotA-E33, MotA-A40, MotA-L81, MotA-Y83, MotA-D128, MotA-E142, MotA-E144, and MotA-R262, have also been previously shown to be important for function (Garza et al., 1996; Togashi et al., 1997; Zhou and Blair, 1997; Hosking and Manson, 2008; Komatsu et al., 2016).

Secondly, this work adds to this list by confirming through point mutation that MotA-Y217N that MotA-Y217 are crucial for function. This finding is similar to the previous report where Deme et al. confirmed the importance of the contact between MotA-Y217 and MotB-W26 via MotB-W26A point mutation (Deme et al., 2020). We further confirmed via point mutations V178I and A179G that MotA-V178 and MotA-A179 were not essential for flagellar function, in contrast to proposals that MotA-V178, MotA-A179 and MotA-A180 residues might control BFM function and ion selectivity (Biquet-Bisquert et al., 2021). None of the point mutations introduced into our non-motile MotA-ASRs were able to restore motility, which could imply these residues were not solely responsible for disruption of motility. However, it must be noted that some of these non-motile ASRs may not be well-folded and we did not test for this.

Thirdly, from the opposite perspective, residues which differed between functional MotA-ASRs and wild-type *E. coli* MotA provided information about variable residues which were not critical for function. This informed the level of general tolerance for significant amino acid substitution across MotA. The presence of multiple distinct (highly variable) residues in the cytoplasmic C-terminal region of MotA, particularly in residues 280–295, indicates that this region is interchangeable, which is similar to previous findings (Muramoto and Macnab, 1998; Santiveri et al., 2020). Although MotA-A187 and MotA-I202 residues are conserved in several proton-powered motors, our functional MotA-ASRs with T187 and V202 indicated that those residues were tolerant to amino acid substitution.

All our motile MotA-ASRs appeared to be proton-powered, that is, they showed no torque-dependence on sodium and were able to rotate in the absence of sodium. Furthermore, growth was inhibited when each of our motile MotA-ASRs was combined with a plug-deleted MotB subunit, indicating that proton leakage was occurring and providing further evidence for proton conductivity in our motile MotA-ASR stator units. It has been proposed that sodium is the ancestral power source for the BFM based on the observation of sodium-dependence in the *Aquifex aeolicus* stator unit (Takekawa et al., 2015). To attempt to answer this using ancestral reconstruction would require well dated phylogenies with appropriate accounting for gene transfer among the stator subunits. We do not address this here, however we note that, to date, none of our ancestral reconstructions of either MotB (Islam et al., 2020) or MotA (this work) have generated a sodium-powered stator unit.

One strength of the ancestral reconstruction approach is that it leverages sequence diversity in contemporary stator proteins. Our results show that even quick, heuristic methods in phylogenetics and ancestral reconstruction can generate functional stator units. Inferred nodes are only estimates, in some cases coarse ones, and may not correctly represent evolutionary history. Improved phylogenetic inferences may improve evolutionary accuracy and the chance of resurrected ancestral stator units being functional. Advanced amino acid substitution models that are selected for higher statistical fit (Minh et al., 2020) should result in phylogenies that are more accurate more deeply in the tree, reducing problems such as long-branch attraction that result from model misspecification (Naser-Khdour et al., 2019; Crotty et al., 2020), and accordingly, more accurate ASRs. Ultimately ASRs may generate non-functional proteins for several reasons, such as failure to fold or even express (Eick et al., 2017). Nevertheless, our results based on functional MotA-ASRs prove successful protein folding through positive function and highlight the role of important regions of MotA that are both conserved and essential for function.

Combinatorial testing of MotA-ASRs from this study with ASRs from our previous MotB-ASR work (Islam et al., 2020) showed that certain MotA-ASRs function only in combination with certain MotB-ASRs. This indicates that there is specificity between MotA and MotB proteins. The promiscuity of stator units (Ito and Takahashi, 2017) and the modular nature of the stator complex (Ridone et al., 2022) may mean that there may be combinations of A and B subunits which are able to power rotation and combinations which cannot. It is therefore suitable to resurrect MotBs that co-existed with MotAs. For this, gene trees of *motA* and *motB* would need to be matched, which should be possible since they are usually syntenic.

Overall, in this work we showcased the utility of ASR for synthesising a range of functional stator units. This delivered insight into the function and evolutionary constraints surrounding an ancient, canonical molecular complex. Future work will examine co-evolutionary history *motA* and *motB* and phylogenies and reconstructions of adjacent genes. This will provide an estimate of the ancestral stator complex while reducing possible complications that may arise from gene transfer.

## Materials and Methods

### Phylogenetic analysis

MotA genes were obtained from select groups of bacterial species and aligned using MAFFT (Katoh et al., 2019) under default settings (MSAs provided in Supplementary Material). Two phylogenies were calculated: the first phylogeny of 178 *motA* homologues was estimated using QuickTree, a neighbour-joining method suitable for large datasets (Howe et al., 2002), with the Kimura translation for pairwise distances. This phylogeny was midpoint-rooted in FigTree (Rambaut, 2014). The second phylogeny was generated using the RAxML-HPC v.8 on XSEDE (Stamatakis, 2014) tool through the CIPRES Science Gateway (Miller et al., 2011) using the PROTGAMMA protein substitution model, LG protein substitution matrix, and a parsimony seed value of 12345. Newick files with full Linnean names and Accession IDs are provided in Supplementary Material.

### Ancestral sequence reconstruction

Both multiple sequence alignments and both Newick files for the two phylogenies in Fig. 1 were used as inputs for CodeML, a maximum likelihood program from the PAML package (Yang, 1997). PAML was run with an empirical model for the amino acid substitution rate using LG as the amino acid ratefile, matching the model used for the RAxML tree generation. As an additional search for functional ancestors, we tested a contemporary ASR method, GRASP (Foley et al., 2022). The 178 QuickTree-generated motA tree (Fig. 1A) was submitted to the GRASP reconstruction webserver with the LG model, and the most probable characters at each site, for each node, was exported for further consideration. Of these reconstructions, two were selected for further synthesis: N41, and N65, corresponding to the nodes 220 and 244 using the labelling system in PAML. All nodes that selected and the ancestral reconstructions that were subsequently synthesised for further testing are indicated in Fig. 1.

### Structural prediction of selected MotA-ASRs

We used ColabFold: AlphaFold2_mmseqs2 notebook version v1.1 (Mirdita et al., 2021) and Alphafold 2.1.1 (Jumper et al., 2021) to predict the 3D structures of our MotA-ASRs and contemporary *E. coli* (RP437) MotAB (MotA_5_MotB_2_) stator complex, respectively. Monomeric structures were generated for individual MotA-ASRs using one copy of ancestral MotA sequence in the default setting. In contrast, an oligomeric structure was generated for the contemporary *E. coli* MotAB complex using five copies of MotA and two copies of MotB (as an oligomeric 5:2 structure) using Alphafold v2.1.1 in multimer mode with the parameters: max template date = 24-1-2022 and prokaryotic_list = true. The resultant structures were aligned with the solved unplugged MotAB stator complex structure of *C. jejuni:* PDB-6YKP (Santiveri et al., 2020) in Pymol (version 2.5.1) for comparison. Monomeric structures were generated for individual MotA-ASRs using one copy of ancestral MotA in the default setting. These monomers were then aligned to individual chains of 6YKP (Supplementary Figure 4).

### Bacterial strains and plasmids and growth conditions

The bacterial strains and plasmids used in this study are shown in full in Supplementary Table 4. The primary strains used in this work are Δ*motA* RP437 (this study) and Δ*motA-ΔmotB* RP437 (this study). All the *E. coli* strains were cultured in LB broth and LB agar [1% (w/v) Bacto tryptone, 0.5% (w/v) Bacto yeast extract, 0.5% (w/v) NaCl and 2% (w/v) Bacto agar for solid media] at 37°C. According to the selective antibiotic resistance pattern of the plasmids, chloramphenicol (CAM), ampicillin (AMP) or kanamycin (KAN) were added to a final concentration of 25 μg/mL, 100 μg/mL and 25 μg/mL, respectively.

### Cloning of selected MotA-ASR sequences

Selected MotA-ASR sequences were ordered as gBlocks from Integrated DNA Technologies (IDT) for cloning into pBAD33 based chimeric plasmid pSHU1234 (Nishino et al., 2015) carrying *pomA* and *potB* (a hybrid of *pomB* and *motB*). The cloning was performed using Circular polymerase extension cloning (CPEC) protocol (Quan and Tian, 2011) with slight modification. Briefly, primers were designed to prepare linearized vector and MotA-ASR inserts that contain overlapping sequences (15-35 bases) between the vector and the inserts. IDT synthesized the designed primers, and the list of all primers is provided in Supplementary Table 5. MotA-ASR inserts and linear vector were prepared by PCR amplification using Q5 high-fidelity (HF) DNA polymerase (NEB). The PCR reaction recipe and condition is provided to supplementary information. The amplified MotA-ASR insert, and linear vector sequences were then separated in a 1% agarose gel and purified using the Qiagen gel purification kit. Finally, the CPEC cloning assembly reaction was performed with the purified MotA-ASR insert and linearized vector using In-Fusion Snap Assembly master mix (Takara) following the manufacturer’s protocol. All the cloned MotA-ASR plasmids were confirmed by colony PCR and sanger sequencing at The Ramaciotti Centre for Genomics (UNSW, Australia) using ASR sequence-specific primers (Supplementary Table 5).

### Δ*motA* strain preparation Cas9-assisted Recombineering

Deletion of *motA* from *E. coli* RP437 was performed using a no-SCAR method, adapted from the previous report (Reisch and Prather, 2015). Briefly, the target strain to be edited (*E. coli RP437*) was sequentially transformed first with the pKD-sgRNA-3’MotA (Sm^+^) plasmid, encoding a sgRNA sequence directed at the 3’ end of motA, and then with the pCas9cr4 (Cm^+^) plasmid to yield a parent strain harboring both plasmids. A slightly modified overlap extension PCR technique (Higuchi et al., 1988) was employed to assemble linear double stranded DNA molecules (dsDNA) using 3 starting dsDNA fragments. The resulting donor DNA was electroporated in the plasmid-bearing host using Gene Pulser Xcell Electroporation Systems (Bio-Rad) and the successfully edited clones were selected via colony PCR and Sanger sequencing of the motAB locus. The list of used primers is provided in Supplementary Table 5.

### Evaluation of functional property of MotA-ASRs

We transformed all the MotA-ASR plasmids into Δ*motA* RP437 and Δ*mot-ΔmotB* RP437 *E. coli* strains following the chemical transformation protocol from NEB for functional characterization. We evaluated the functional property of resurrected MotA-ASRs using swim plate motility assay and free-swimming assay following the previous protocol (Morimoto et al., 2017; Islam et al., 2020). Briefly, for swim plate motility assay, we inoculated fresh single colonies of ten MotA-ASR transformed Δ*motA* and Δ*mot-ΔmotB E. coli* on (0.02%) arabinose and chloramphenicol (25 μg/ml) containing semi solid LB swim agar (0.25%) with a sterile toothpick and incubated at 30□ for 14h to allow proper development of a swimming ring. Swimming zones were visually checked, imaged with the Chemi Doc MP Imaging System (Bio-Rad).

For free swimming assay, the overnight culture of the MotA-ASR transformed Δ*motA E. coli* strains were subcultured with a 50-fold dilution into fresh TB broth and incubated for 5 h with 180 rpm at 30°C. At OD600 ~0.80, the cells were washed once with motility buffer (10 mM potassium-phosphate, 10 mM lactic acid, 100 mM NaCl, and 0.1 mM EDTA, pH 7.0). Then, cell suspensions were added to the tunnel slide and the swimming speed of the cells was observed using phase-contrast microscopy and a 20 s video was recorded at 20 frames per second (fps) through a 20x objective (Nikon) with a camera (Chameleon3 CM3, Point Grey Research). Finally, the swimming speed of the cells in was calculated using LabVIEW 2019 software (National Instruments).

### Determination and confirmation of the power source of MotA-ASRs

We used a minimal swim plate in low Na^+^ condition (~1 mM) without tryptone or yeast extract and where NaCl had been substituted with KCl (Islam et al., 2020; Ridone et al., 2022), to determine which one between sodium (Na^+^) and proton (H^+^) was the power source of our MotA-ASRs. The minimal swim plates were inoculated with a fresh single colony of MotA-ASR transformed Δ*motA E. coli* with a sterile toothpick and incubated at 30 for 16h. After incubation, we checked whether the tested MotA-ASRs could produce any swimming ring or not in the absence of Na^+^ and compared the phenotype with the control Na^+^ (*pomApotB*) and H^+^ (*motAmotB*) swimmers.

We further used the tethered cell assay to confirm the H^+^ dependent phenotype of the previous minimal swim plate assay following a modified version of earlier protocol (Nishiyama and Kojima, 2012). Briefly, the overnight culture of the MotA-ASR transformed Δ*motA E. coli* strains were subcultured with a 50-fold dilution into fresh TB broth and incubated for 5 h with 180 rpm at 30°C. At OD600 ~0.80, the flagella of the cells were sheared by passing the culture multiple times (~35) through a 26G needle syringe. After shearing the flagella, the cells were washed three times with motility buffer. Cells were then attached on glass slides pre-treated with an anti-*E. coli* flagellin antibody with a 1:10 dilution (Nishiyama and Kojima, 2012) and washed sequentially with a Na^+^ concentration gradient containing motility buffer. The rotational speed of the cells was observed using phase-contrast microscopy (Nikon) and was recorded at 20 frames per second (FPS) through the 40X objective with a camera (Chameleon3 CM3, Point Grey Research). Rotational speed of the cells was analysed and calculated using Lab view 2019 software (National Instruments) and rotational speeds were calculated from 20 individual cells.

### Growth curve assays

Growth curves of MotA-ASRs were determined using a slightly modified protocol as described (Islam et al., 2020). Briefly, overnight cultures of the test strains with OD600 1.0 were diluted 1:100 in 96-well plates (Corning) with fresh LB broth, 0.02% (w/v) arabinose and 25 μg/ml chloramphenicol and incubated at 37 □. The OD600 was measured every hour in a microplate reader (FLUOstar OPTIMA, BMB LABTECH) with a brief shaking interval before each measurement. The experiment was performed in a triplicate.

### Point mutations in WT MotA, MotA-ASRs and plug deletion from MotB

We performed the selected point mutation to test the roles of our identified conserved residues for the disruption or rescue of motility in *E. coli* and functional MotA-ASRs through *QuickChange Site-Directed Mutagenesis Kit* (Agilent). Initially, mutagenic oligonucleotide primers were designed in Agilent quick change primer design software at https://www.agilent.com/store/primerDesignProgram.jsp (Supplementary Table 5.). Then the mutant strands were synthesized following the Agilent recommended protocol by PCR amplification. Finally, the desired point mutants were confirmed by Sanger sequencing before functional analysis. We constructed plug deleted pMotB by deleting residues 51-70 from *motB* using Site-directed, Ligase-Independent Mutagenesis (SLIM), following previous protocol (Chiu et al., 2008).

## Acknowledgements

The authors acknowledge use of facilities in the Structural Biology Facility within the Mark Wainwright Analytical Centre– UNSW, funded in part by the Australian Research Council Linkage Infrastructure, Equipment and Facilities Grant: ARC LIEF 190100165. MABB acknowledges funding support from a Scientia Fellowship from UNSW, a CSIRO Future Science Platform in Synthetic Biology Project Grant, and Australian Research Council Discovery Project DP190100497 and Human Frontiers Science Program Project Grant RGY0072/21.

