## Supplementary Figures 1-12, Supplementary Tables 1-3. for "Ancestral reconstruction of the MotA stator subunit reveals that conserved residues far from the pore are required to drive flagellar motility"

### Supplementary Material

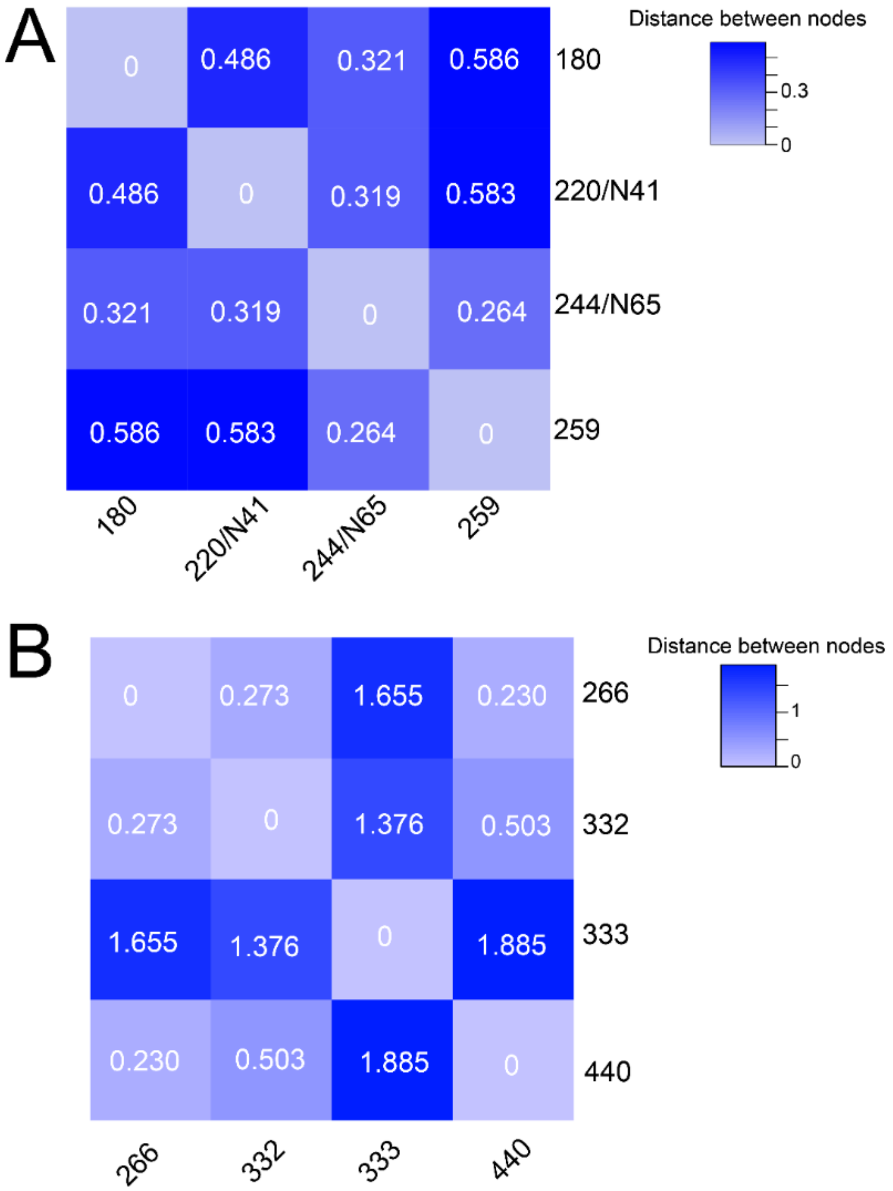

**Supplementary Figure 1:** Pairwise distances between all selected nodes. (A) Pairwise distances of the nodes (180, 220/N41, 244/N65, and 259) were calculated for the phylogenetic tree of 178 MotA homologs (main Fig. 1A) from the total branch distance of the shortest connection between two nodes. (B) Pairwise distances of the nodes (266, 332, 333, and 440) were likewise calculated from the branch lengths of the phylogenetic tree of 264 MotA homologs (main Fig. 1B). Branch length unit was substitutions per site. The heatmaps were generated in the heatmap2 program from the Galaxy Australia website (https://usegalaxy.org.au/ ).

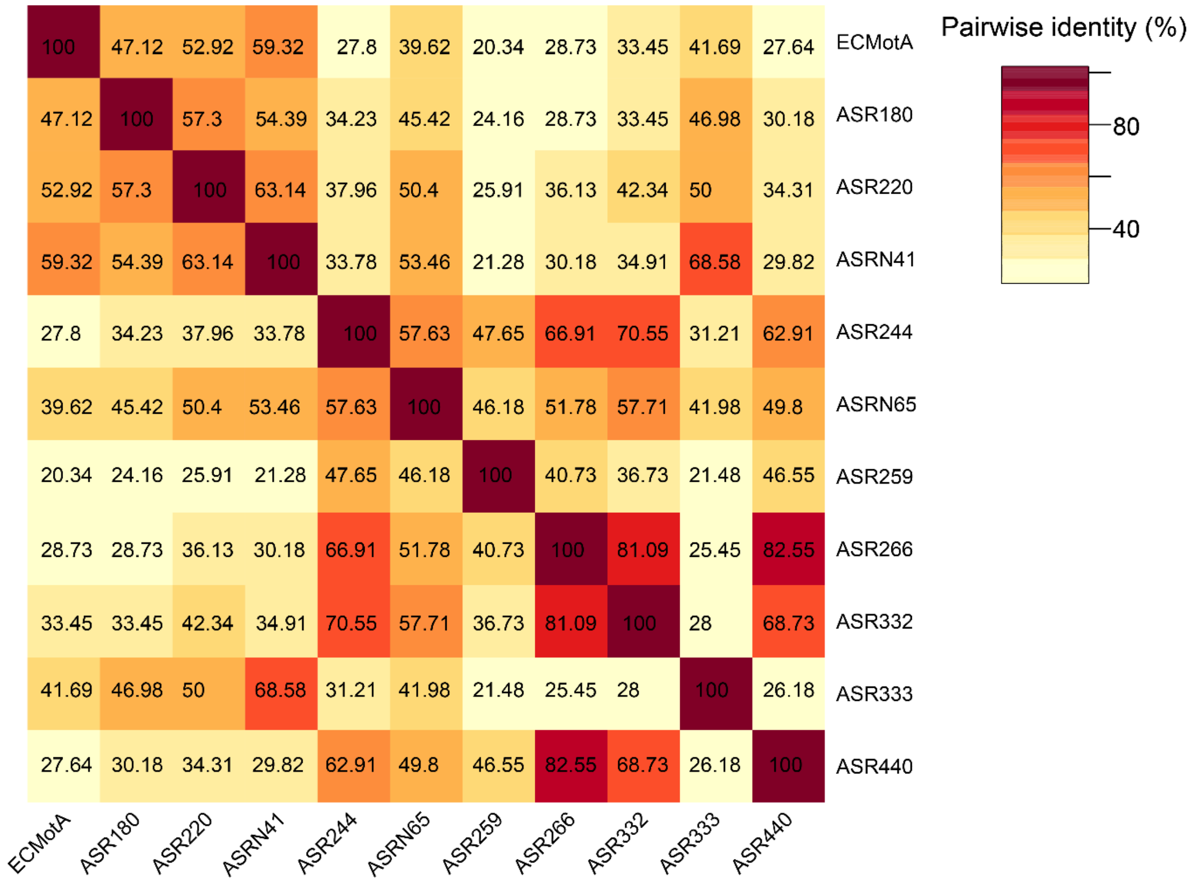

**Supplementary Figure 2:** Pairwise percentage similarity (identity) displayed as matrix of all ten selected ASRs alongside wild-type *E. coli* MotA. The identity matrix was calculated using Clustal Omega (https://www.ebi.ac.uk/Tools/msa/clustalo/) and the heatmap was generated in the heatmap2 program from the Galaxy Australia website (https://usegalaxy.org.au/ ).

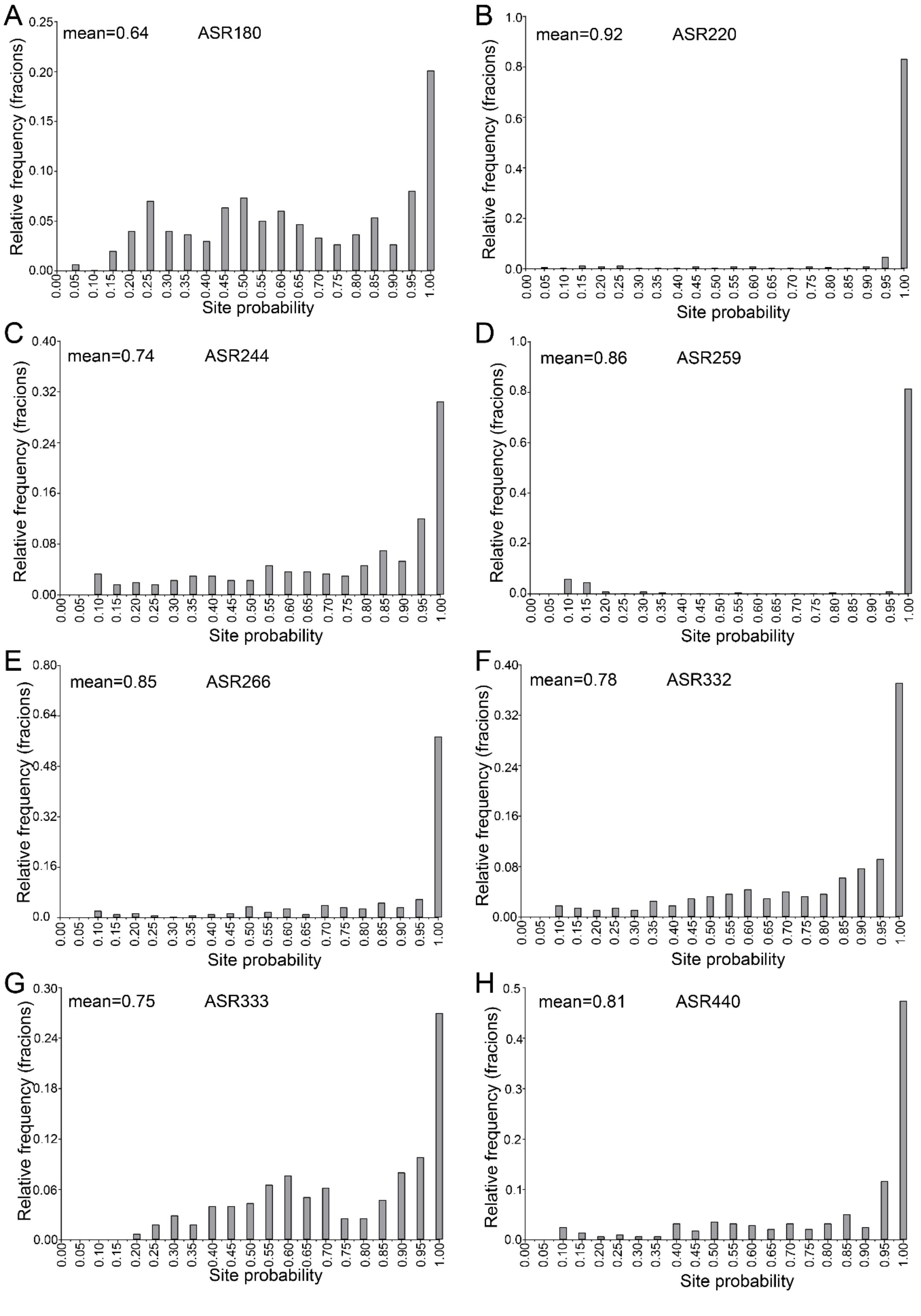

**Supplementary Figure 3:** Histogram of posterior probabilities from the PAML ancestral reconstruction for most common amino acid at each site, for each ASR, at the 8 selected nodes. Mean probability across all sites is shown inset in top left.

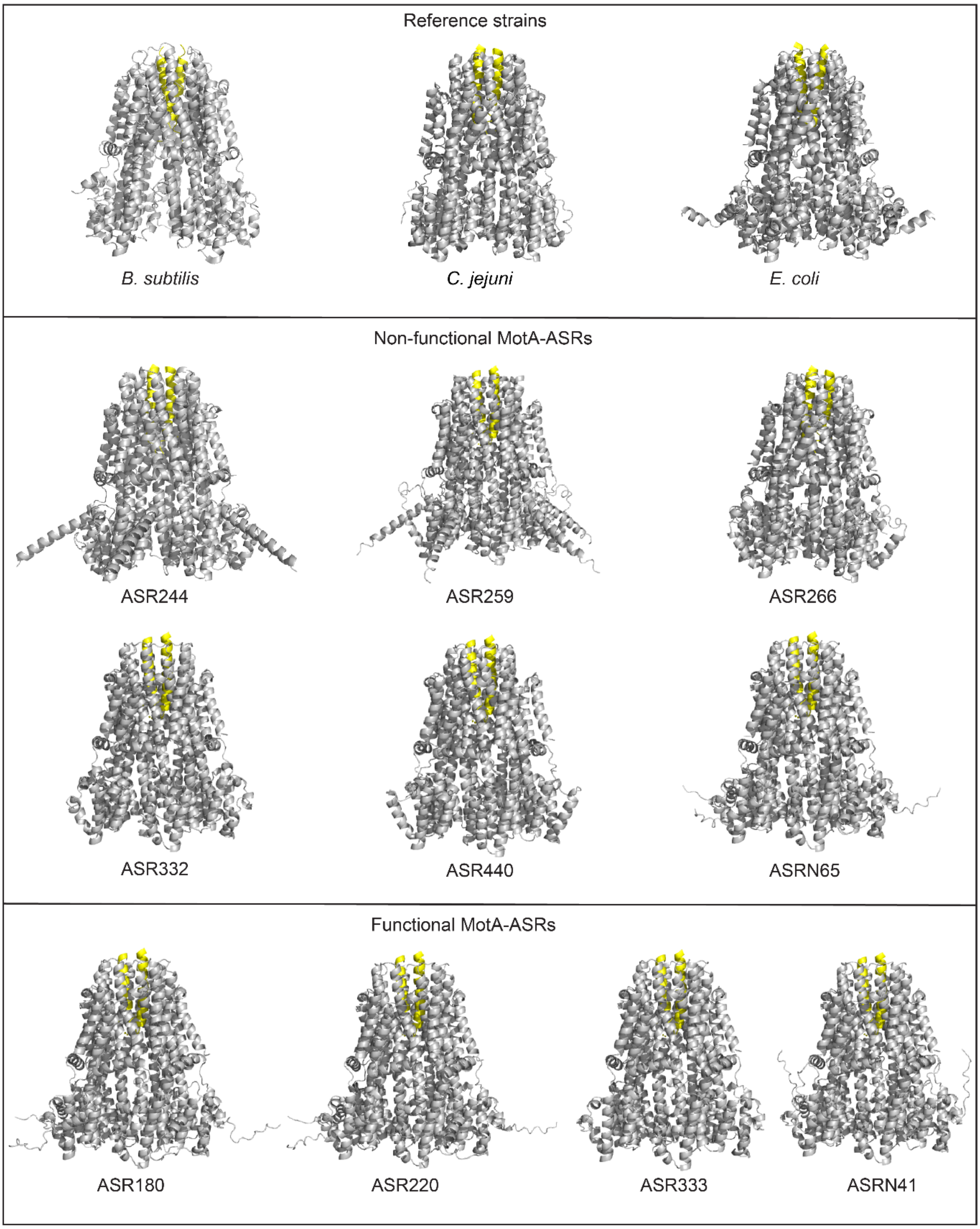

**Supplementary Figure 4:** ColabFold structural models for MotA-ASRs. Monomer subunits for *E. coli* MotA and each of the ten MotA-ASRs were aligned with pentamer MotA from *C. jejuni* $MotA_5MotB_2$ stator complex structure: PDB-6YKP (Santiveri et al., 2020). MotA pentamers and MotB dimers are show in grey and yellow, respectively. *B. subtilis* structure: PDB-6YSL (Deme et al., 2020), *E. coli* structure from Alphafold model in this work.

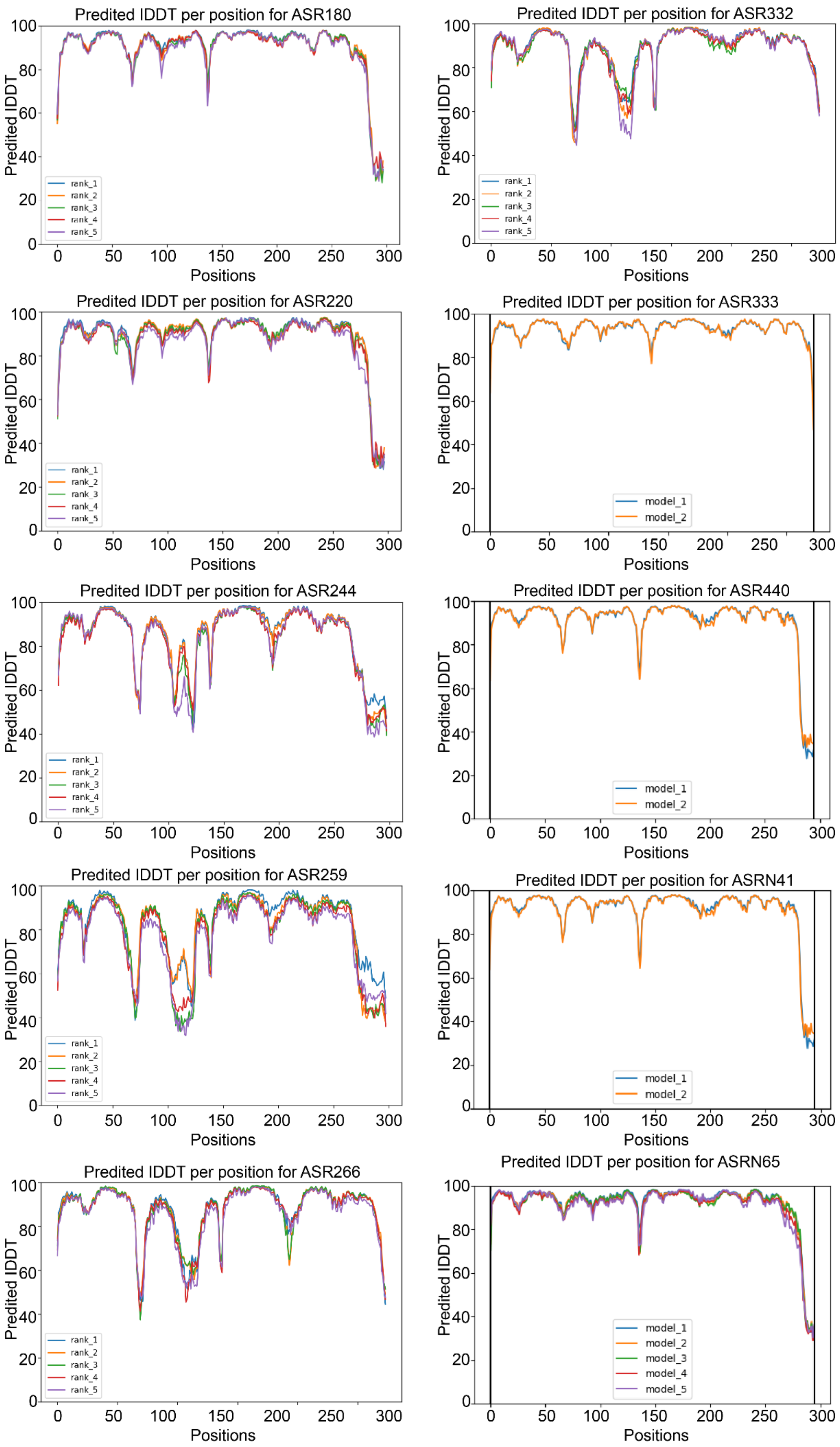
Predited IDDT per position for ASR180
Predited IDDT per position for ASR332
Predited IDDT per position for ASR220
Predited IDDT per position for ASR333
Predited IDDT per position for ASR244
Predited IDDT per position for ASR440
Predited IDDT per position for ASR259
Predited IDDT per position for ASRN41
Predited IDDT per position for ASR266
Predited IDDT per position for ASRN65
Predited IDDT
Positions
rank_1
rank_2
rank_3
rank_4
rank_5
model_1
model_2
model_3
model_4
model_5

27

**Supplementary Figure 5:** Predicted LDDT per position for the Colabfold models of MotA-ASRs. All the plots of the MotA-ASRs show the predicted LDDT per position for the 2-5 models obtained from Colabfold output. The top-ranked models were selected for each MotA-ASR and aligned with the *C. jejuni* MotAB complex (PDB:6YKP) shown in Supplementary Figure 4.

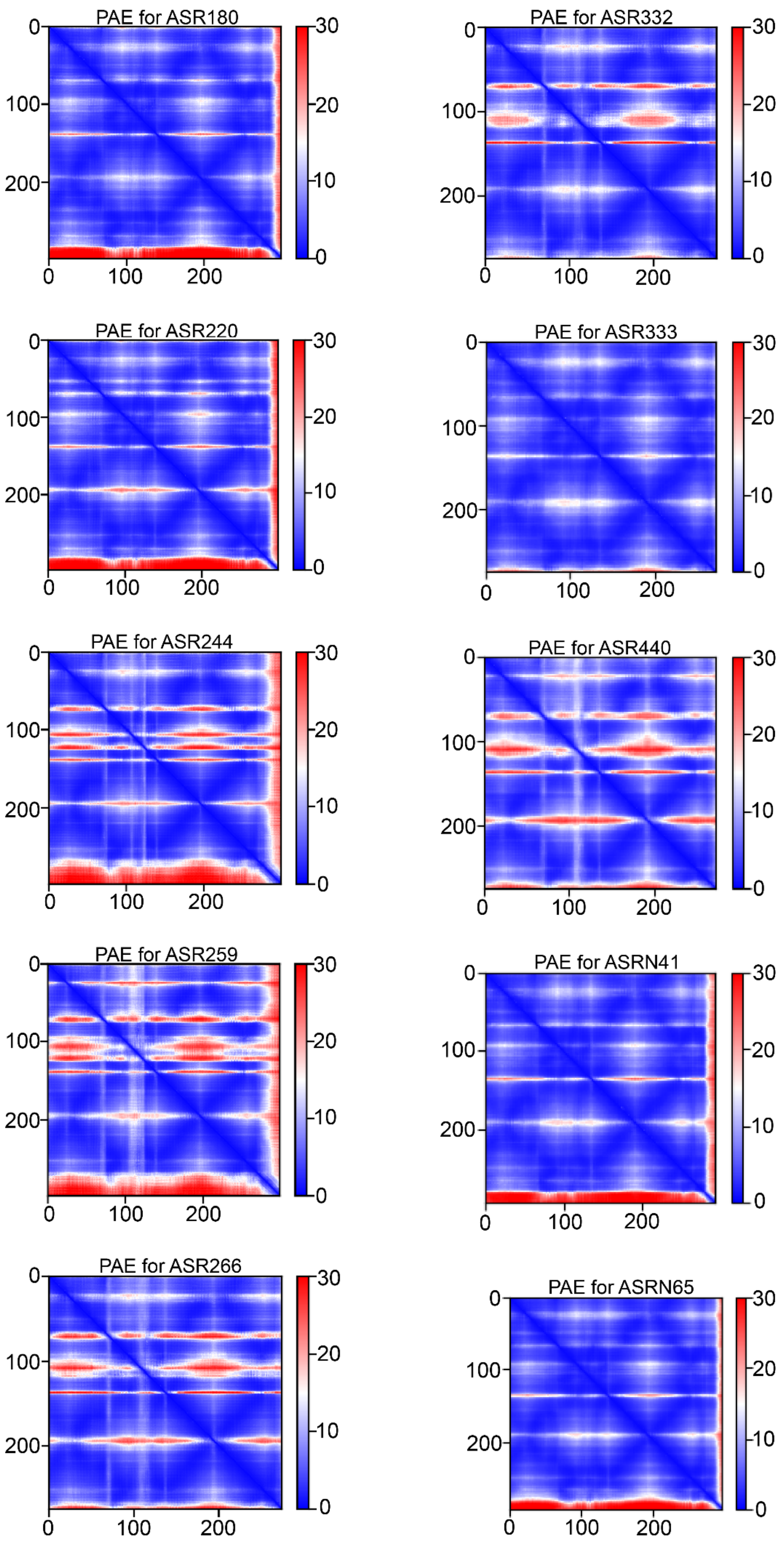
PAE for ASR180
PAE for ASR332
PAE for ASR220
PAE for ASR333
PAE for ASR244
PAE for ASR440
PAE for ASR259
PAE for ASRN41
PAE for ASR266
PAE for ASRN65

**Supplementary Figure 6:** Predicted aligned errors (PAE) for the MotA-ASR Colabfold models showed in Supplementary Figure 4.

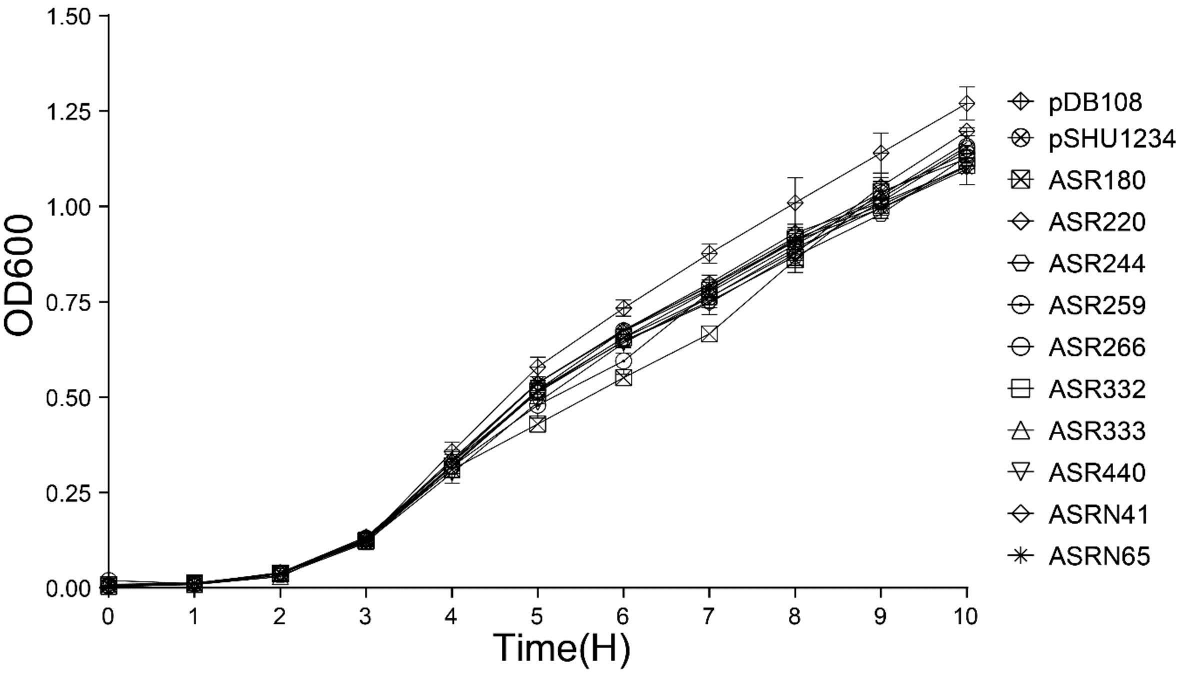

**Supplementary Figure 7:** Growth curve of MotA-ASRs in the presence of WT *E. coli* MotB. Ten MotA-ASRs transformed into a Δ*motA* RP437 strain and the controls, pSHU1234 (*pomApotB*) and pDB108 (*motAmotB*) transformed into a Δ*motAmotB* RP437. All of the transformed cells were grown in LB broth with appropriate antibiotic selection and 0.2% arabinose in a 96-well microtiter plate, and the OD was measured hourly.

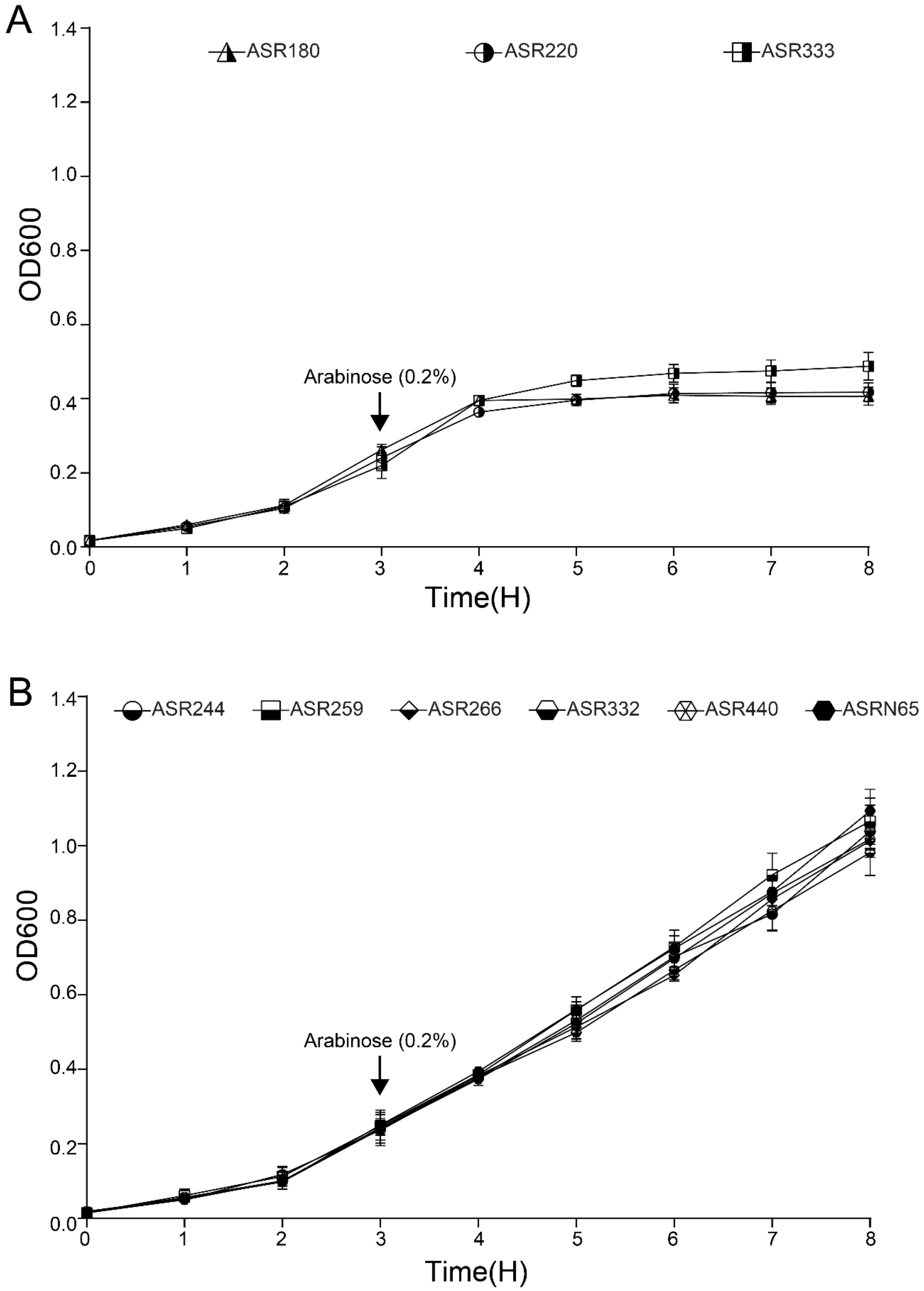

**Supplementary Figure 8:** Growth curve of MotA-ASRs in the presence of plug deleted (Δ51-70) WT *E. coli* MotB. (A) and (B) are showing the growth curve of the functional MotA-ASRs (ASR180, ASR220 and ASR33) and non-functional MotA-ASRs (ASR244, ASR259, ASR266, ASR332, ASR440 and ASRN65), respectively. Both the functional and non-functional MptA-ASRs were co-transformed with MotB$_{\Delta 51\text{-}70}$ into a Δ*motAmotB* RP437 strain and grown in LB broth with CAM and AMP antibiotics in a 96-well microtiter plate. 0.2% arabinose was added after three hours of incubation.

```
ECMotA 1 --MLILLGYLVVLGTVFGGYLMTGGSLGALYQPAELVIIAGAGIGSFIVGNNGKAIKGTL 58
ASR180 1 MDMQKIIGIVIIFGCVFGGYLMAGGKLDVIWQPAELMIIGGAGVGAFIIGNPLTVIKETA 60
ASR220 1 MDMAKIIGIIVVFASVLGGYVLSHGKIAALIQPFEVLIIGGAAFGAFLQANPGHMTMHVI 60
ASR333 1 --MFAIIGIIVVFACVFGGFLVAGGHLGVLWQPFELLIIGGAALGAFIISNPAKVLKATG 58
ASRN41 1 --MLVIIGIIVVLGSVLGGYVLSHGKLAALIQPAELLIIGGAAIGAFLVANPGKVIKATV 58
           *  ::* ::::. *:**:::  * : .: ** *::**.**..*:*: .*       .
ECMotA 59 KALPLLFRRSKYTKAMYMDLLALLYRLMAKSRQMGMFSLERDIENPRESEIFASYPRILA 118
ASR180 61 NGLGKVFKGPKWKEEHYRDLLALLYELMKTVRSKGLIALEEHIENPQESSIFNRYPKVLK 120
ASR220 61 KKSMKMFGGSRFSHAYYLEVLGLVYEILNKSRREGMMAIEADIEDPAASPIFAKYPTVLK 120
ASR333 59 KALAKVFKGSKYKKEDYLELLALLYELFQTARKEGPMALEKHIEDPHESPIFQQYPKFLK 118
ASRN41 59 KGLMKVFRGSKYSKADYLDLLALLYELLNKSRREGMMAIEADIENPAESPIFSKYPKILA 118
          :    :*   ::..  * ::*.*:*.:: . *  * :::* .**:*  * **  ** .*
ECMotA 119 DSVMLDFIVDYLRLIISGHMNTFEIEALMDEEIETHESEAEVPANSLALVGDSLPAFGIV 178
ASR180 121 DHQLVSFICDNLRLMVMGNMDPHEIEGIMEQEIEAIEEDLLKPSHALQSMGDALPAFGIV 180
ASR220 121 DERMTAFICDYLRIMSSGNMAPHELEGLFDMELLSMKEELEHPSHAVTGIADGMPGFGIV 180
ASR333 119 DHHAVHFLCDTLRLIVSGSMNPHEVEDLMDEEIETHHHEQHQPAHAIQTVADGLPALGIV 178
ASRN41 119 DHHLVDFICDYLRMMVSGNMDPHEIEGLMDMEIETMHHEAEVPAHALTKVADGLPGFGIV 178
           *     *: * **::  * *  .*:* ::: *: : . :   *::::  :.*.:*.:***
ECMotA 179 AAVMGVVHALGSADRPAAELGALIAHAMVGTFLGILLAYGFISPLATVLRQKSAETSKMM 238
ASR180 181 AAVLGIIKTMGSIDESPAVIGAKIAAALVGTFLGVFMAYGLLGPLATRLEAQVEKEGALY 240
ASR220 181 AAVLGIVVTMASLGGDQAAIGMHVGAALVGTFFGILAAYGFFGPLATSLEHDAKEELNLY 240
ASR333 179 AAVLGVVKTMGSLNEPPEKLGHLIASALVGTFLGVFLAYGFVGPLATKLKQKVDEEAKYF 238
ASRN41 179 AAVLGVVITMGSLGGPQEEIGHHVGAALVGTFLGILLAYGFVGPLATSLEHRAEEETKMY 238
           ***:*:: ::.* .     :*  :. *:****:*:: ***:..**** *.    :
ECMotA 239 QCVKVTLLSNLNGYAPPIAVEFGRKTLYSSERPSFIELEEHVRAVKNPQQQTTTEEA- 295
ASR180 241 KIVKAVLVAHLHGNAPQIAVEAGRKTIPSDHRPSFAELEEALTEQPGEGAGKAAPKAA 298
ASR220 241 EAIKASLVASASGMPPSLAVEFGRKVLYPKHRPSFAELEQAVRGRKSAAPGAAGSEAA 298
ASR333 239 HCIKAALLALQHGYPPQVCVEYARKALYPEERPSFE---------------------- 274
ASRN41 239 QAIKVALVASVNGYPPQLAVEFGRKALPSNVRPSFAELEEAVRGRKAPASQATEEEAE 296
           . :*. *::   *  * :.** .**.:  . ****
```

**Supplementary Figure 9:** Multiple sequence alignment of WT *E. coli* MotA and the four functional MotA-ASRs (ASR180, ASR220, ASR333 and ASRN41). Below each site (i.e., position) of the protein sequence alignment is a key denoting conserved sites (*), sites with conservative replacements (:), sites with semi-conservative replacements (.), and sites with non-conservative replacements ( ).

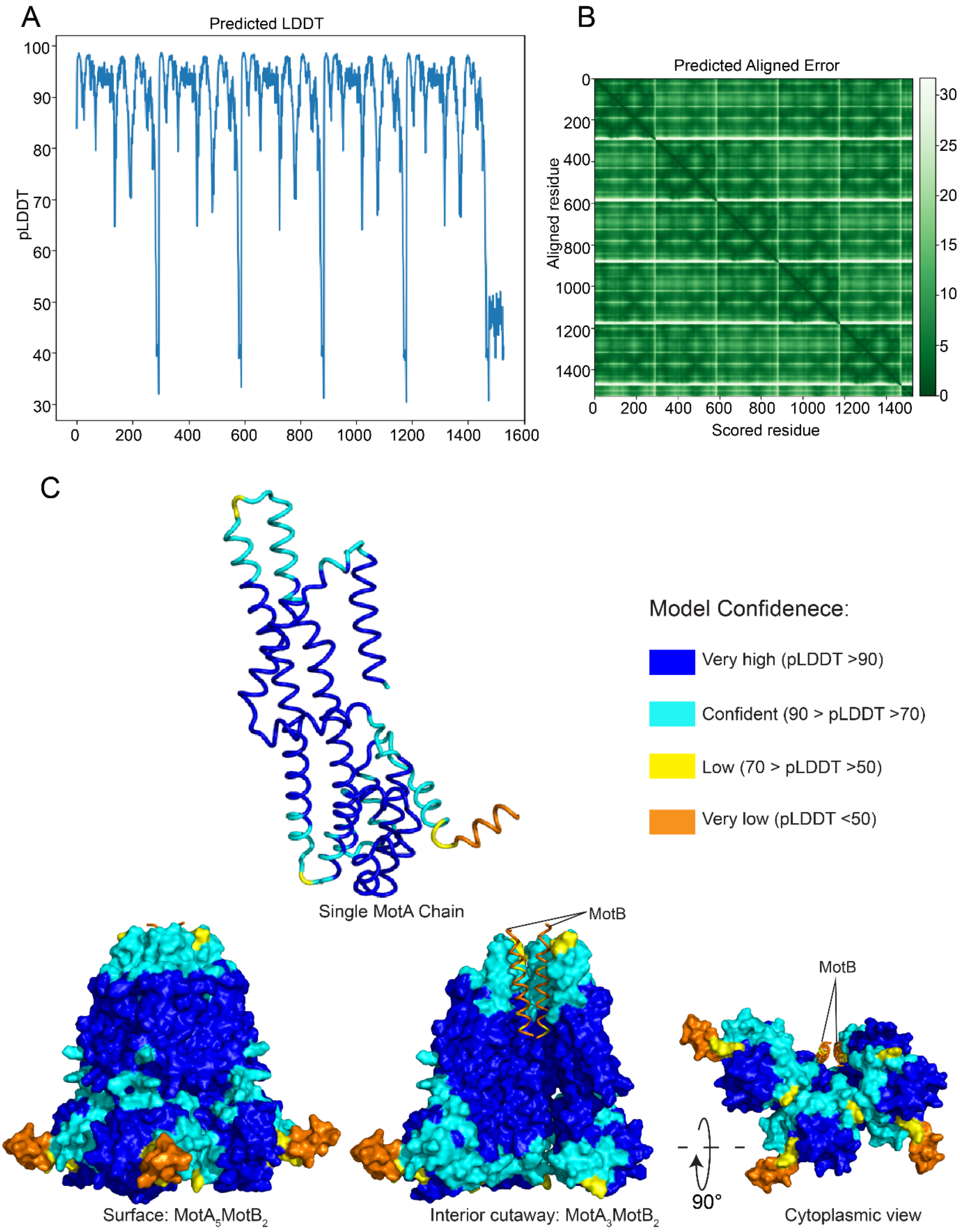

**Supplementary Figure 10:** Alphafold structure prediction of WT *E. coli* MotAB complex (5:2). (A) Predicted LDDT per position for five copies of MotA (295 residues) and two copies of MotB (26 residues, TM domain). (B) Predicted aligned error (PAE) for the $MotA_5MotB_2$ model. (C) *E. coli* MotAB Alphafold model colour-coded by the pLDDT confidence scores.

The confidence of the model is shown on a single MotA chain in the cartoon representation (upper left) as well as on the surface (lower left) and the interior of the MotAB complex (lower middle and lower right).

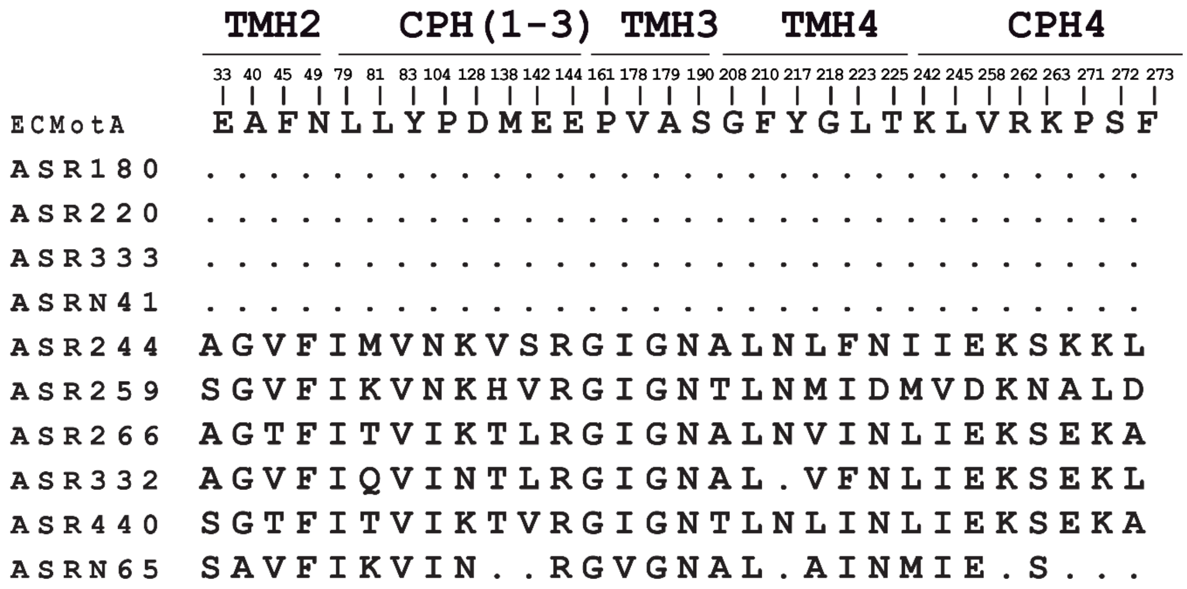

**Supplementary Figure 11:** 30 proposed critical residues identified from sequence comparison between WT *E. coli* MotA, and both functional and non-functional MotA-ASRs. Sites were identified by those sites which were conserved across functional MotA-ASRs but different in non-functional MotA-ASRs. Location on the MotA regions are are labelled on top: transmembrane helix 2 (TMH2), cytoplasmic helix 1-3 (CPH1-3), transmembrane helix 3 (TMH3), transmembrane helix 4 (TMH4) and cytoplasmic helix 4 (CPH4). Identical residues are presented with dot (.).

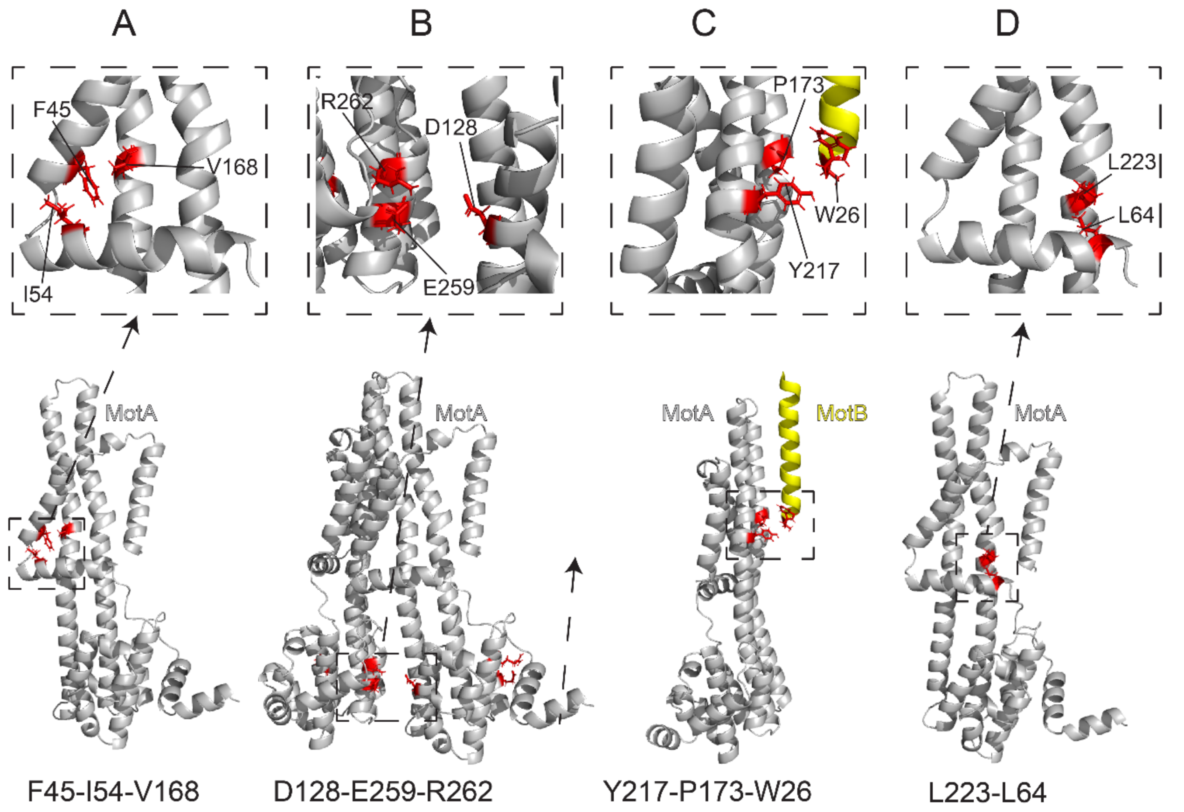

**Supplementary Figure 12:** Predicted molecular contacts of critical residues. Grey color represents MotA and Yellow color represents MotB. (A) Interaction of MotA-F45 with MotA-I54 and MotA-V168. (B) Interaction of MotA-D128 with other MotA-E259 and R262. (C) Interaction of MotA-Y217 with MotA-P173 and MotB-W26. (D) Interaction of MotA-L223 with MotA-L64.

**Supplementary Table 1:**

RMSD for each chain of *E. coli* MotAB Alphafold model in comparison with each of 6YKP, 6YKM and 6YSL.

| *E. coli* MotAB | 6YKP | RMSD |
|---|---|---|
| Chain A (MotA1) | Chain A (MotA1) | 2.157 |
| Chain B (MotA2) | Chain B (MotA2) | 2.14 |
| Chain C (MotA3) | Chain C (MotA3) | 2.263 |
| Chain D (MotA4) | Chain D (MotA4) | 2.008 |
| Chain E (MotA5) | Chain E (MotA5) | 2.337 |
| Chain F (MotB1) | Chain F (MotB1) | 3.517 |
| Chain G (MotB2) | Chain G (MotB2) | 3.51 |
| *E. coli* MotAB | 6YKM | RMSD |
| Chain A (MotA1) | Chain A (MotA1) | 2.251 |
| Chain B (MotA2) | Chain B (MotA2) | 2.19 |
| Chain C (MotA3) | Chain C (MotA3) | 2.299 |
| Chain D (MotA4) | Chain D (MotA4) | 2.068 |
| Chain E (MotA5) | Chain E (MotA5) | 2.306 |
| Chain F (MotB1) | Chain F (MotB1) | 3.516 |
| Chain G (MotB2) | Chain G (MotB2) | 3.424 |
| *E. coli* MotAB | 6YSL | RMSD |
| Chain A (MotA1) | Chain C (MotA1) | 1.926 |
| Chain B (MotA2) | Chain D (MotA2) | 2.651 |
| Chain C (MotA3) | Chain E (MotA3) | 2.077 |
| Chain D (MotA4) | Chain F (MotA4) | 2.745 |
| Chain E (MotA5) | Chain G (MotA5) | 2.233 |
| Chain F (MotB1) | Chain A (MotB1) | 3.386 |
| Chain G (MotB2) | Chain B (MotB2) | 3.178 |

**Supplementary Table 2:**

List of molecular contacts of the identified 30 conserved MotA residues determined in the alphafold structure of *E. coli* MotA/MotB stator complex measured using pymol (V4.6.0). Inter-subunit contacts are shown in bold.

| Conserved residue | Interacting residue | Estimated distance between the residues showed in angstrom (Å) |
|---|---|---|
| MotA-E33 | MotA-G208 | 2.9 |
| MotA-E33 | MotA-A179 | 2.4 |
| **MotA-A40** | **MotA-V14** | 2.5 |
| MotA-F45 | MotA-I54 | 2.8 |
| MotA-F45 | MotA-V168 | 2.5 |
| MotA-N49 | MotA-L167 | 4.3 |
| MotA-L79 | MotA-V258 | 2.8 |
| MotA-L81 | MotA-I109 | 2.3 |
| MotA-Y83 | MotA-P255 | 2.8 |
| MotA-P104 | MotA-V127 | 3 |
| **MotA-D128** | **MotA-E259** | 7.1 |
| **MotA-D128** | **MotA-E262** | 4.9 |
| **MotA-M138** | **MotA-Y252** | 3.3 |
| MotA-E142 | MotA-L246 | 2.8 |
| MotA-E144 | MotA-V243 | 5.8 |
| MotA-P161 | MotA-A60 | 3 |
| **MotA-V178** | **MotA-F210** | 2.4 |
| MotA-A179 | MotA-G208 | 2.5 |
| MotA-S190 | MotA-L198 | 3.2 |
| MotA-Y217 | MotA-P173 | 3 |

| | | |
|---|---|---|
| MotA-Y217 | MotB-W26 | 4.1 |
| MotA-G218 | MotA-V10 | 2.4 |
| MotA-L223 | MotA-L64 | 2.7 |
| MotA-L223 | MotA-L165 | 3.7 |
| **MotA-T225** | **MotA-G48** | 3.3 |
| MotA-K242 | MotA-D148 | 2.6 |
| MotA-L245 | MotA-Y129 | 3.2 |
| MotA-L245 | MotA-M147 | 2.3 |
| **MotA-K263** | **MotA-Y129** | 3.9 |
| MotA-P271 | MotA-R262 | 2.8 |
| MotA-F273 | MotA-E259 | 2.5 |

**Supplementary Table 3:**

List of the point mutants and their motility status that we generated in this study. 6PMA = G40A + G40A + I178V + G179A + N217Y + L218G + E262R

| Point mutant name | Motility status | Swimming ring production |
|---|---|---|
| *E. coli* MotA-V178I | Motile | + |
| *E. coli* MotA-A179G | Motile | + |
| *E. coli* MotA-Y217N | Non-motile | - |
| ASR180-V178I | Motile | + |
| ASR180-A179G | Motile | + |
| ASR180-Y217N | Non-motile | - |
| ASR220-V178I | Motile | + |
| ASR220-A179G | Motile | + |
| ASR220-Y217N | Non-motile | - |
| ASR333-V178I | Motile | + |
| ASR333-A179G | Motile | + |
| ASR333-Y217N | Non-motile | - |
| ASRN41-V178I | Motile | + |
| ASRN41-A179G | Motile | + |
| ASRN41-Y217N | Non-motile | - |
| ASR244-G40A | Non-motile | - |
| ASR244-I178V | Non-motile | - |
| ASR244-G179A | Non-motile | - |
| ASR244-N217Y | Non-motile | - |
| ASR244-L218G | Non-motile | - |
| ASR244-E262R | Non-motile | - |

| | | |
|---|---|---|
| ASR244-6PMA | Non-motile | - |
| ASR259-G40A | Non-motile | - |
| ASR259-I178V | Non-motile | - |
| ASR259-G179A | Non-motile | - |
| ASR259-N217Y | Non-motile | - |
| ASR259-M218G | Non-motile | - |
| ASR259-E262R | Non-motile | - |
| ASR259-6PMA | Non-motile | - |
| ASR266-G40A | Non-motile | - |
| ASR266-I178V | Non-motile | - |
| ASR266-G179A | Non-motile | - |
| ASR266-N217Y | Non-motile | - |
| ASR266-V218G | Non-motile | - |
| ASR266-E262R | Non-motile | - |
| ASR266-6PMA | Non-motile | - |
| ASR332-G40A | Non-motile | - |
| ASR332-I178V | Non-motile | - |
| ASR332-G179A | Non-motile | - |
| ASR332-N217Y | Non-motile | - |
| ASR332-V218G | Non-motile | - |
| ASR332-E262R | Non-motile | - |
| ASR332-6PMA | Non-motile | - |
| ASR440-G40A | Non-motile | - |
| ASR440-I178V | Non-motile | - |
| ASR440-G179A | Non-motile | - |

| | | |
|---|---|---|
| ASR440-N217Y | Non-motile | - |
| ASR440-L218G | Non-motile | - |
| ASR440-E262R | Non-motile | - |
| ASR440-6PMA | Non-motile | - |
| ASRN65-G40A | Non-motile | - |
| ASRN65-I178V | Non-motile | - |
| ASRN65-G179A | Non-motile | - |
| ASRN65-N217Y | Non-motile | - |
| ASRN65-A218G | Non-motile | - |
| ASRM65-E262R | Non-motile | - |
| ASRM65-6PMA | Non-motile | - |

**Supplementary Table 4:**

List of strains and plasmids

| **Strain** | **Description** | **Reference** |
|---|---|---|
| RP437-ΔMotA/MotB | *E. coli* (ΔMotA, ΔMotB) | This study |
| RP3087-ΔMotA | *E. coli* (ΔMotA) | |
| **Plasmids** | **Description** | **Reference** |
| pSHU1234 | PomA and PotB, Ara, $\text{CAM}^{R}$ | Kojima et al., 2008 |
| pBAD33 | Empty vector, $\text{CAM}^{R}$ | Guzman et al., 1995 |
| pDB108 | MotA and MotB, $\text{CAM}^{R}$ | David F Blair |
| pMotB | pDB108 ΔMotA $\text{CAM}^{R}$ | Islam et al., 2020 |
| pMotB | pDFB27 ΔMotA $\text{AMP}^{R}$ | David F Blair |
| pNT8 | pSBETa- $\text{motB}_1^{Aa}$, $\text{KAN}^{R}$ | Takekawa et al., 2015 |
| pNT9 | pSBETa- $\text{motB}_2^{Aa}$, $\text{KAN}^{R}$ | Takekawa et al., 2015 |
| P180 | PotB and MotA-ASR180, pSHU1234 backbone, $\text{CAM}^{R}$ | This study |
| P220 | PotB and MotA-ASR220, pSHU1234 backbone, $\text{CAM}^{R}$ | This study |

| | | |
|---|---|---|
| P244 | PotB and MotA-ASR244, pSHU1234 backbone, CAM$^R$ | This study |
| P259 | PotB and MotA-ASR259, pSHU1234 backbone, CAM$^R$ | This study |
| P266 | PotB and MotA-ASR266, pSHU1234 backbone, CAM$^R$ | This study |
| P332 | PotB and MotA-ASR332, pSHU1234 backbone, CAM$^R$ | This study |
| P333 | PotB and MotA-ASR333, pSHU1234 backbone, CAM$^R$ | This study |
| P440 | PotB and MotA-ASR440, pSHU1234 backbone, CAM$^R$ | This study |
| pN41 | PotB and MotA-ASRN41, pSHU1234 backbone, CAM$^R$ | This study |
| pN65 | PotB and MotA-ASRN65, pSHU1234 backbone, CAM$^R$ | This study |
| p758 | MotB-ASR758, pSHU1234 backbone, CAM$^R$ | Islam et al., 2020 |
| p759 | MotB-ASR759, pSHU1234 backbone, CAM$^R$ | Islam et al., 2020 |
| p760 | MotB-ASR760, pSHU1234 backbone, CAM$^R$ | Islam et al., 2020 |
| p765 | MotB-ASR765, pSHU1234 backbone, CAM$^R$ | Islam et al., 2020 |
| p908 | MotB-ASR908, pSHU1234 backbone, CAM$^R$ | Islam et al., 2020 |
| p981 | MotB-ASR981, pSHU1234 backbone, CAM$^R$ | Islam et al., 2020 |
| p1024 | MotB-ASR1024, pSHU1234 backbone, CAM$^R$ | Islam et al., 2020 |
| P1170 | MotB-ASR1170, pSHU1234 backbone, CAM$^R$ | Islam et al., 2020 |
| p1239 | MotB-ASR1024, pSHU1234 backbone, CAM$^R$ | Islam et al., 2020 |
| p1246 | MotB-ASR1246, pSHU1234 backbone, CAM$^R$ | Islam et al., 2020 |
| p1457 | MotB-ASR1457, pSHU1234 backbone, CAM$^R$ | Islam et al., 2020 |
| p1459 | MotB-ASR1459, pSHU1234 backbone, CAM$^R$ | Islam et al., 2020 |
| p1501 | MotB-ASR1501, pSHU1234 backbone, CAM$^R$ | Islam et al., 2020 |

**Supplementary Table 5:**

List of primer sequences and PCR conditions used in this work.

| Primer name | Primer sequence |
|---|---|
| 180 GBLOCK - FW | GGAGTGCTTTATGGATATGCAGAAAATTATTGGTATC |
| 180 PSHU - RV | CTGCATATCCATAAAGCACTCCTCACGC |
| 180 PSHU - FW | GAAAGCAGCATAACTTGGAGAATTCATATGGATGAT |
| 180 GBLOCK - RV | ATTCTCCAAGTTATGCTGCTTTCGGGG |
| 220 GBLOCK - FW | AGTGCTTTATGGACATGGCTAAAATCATCG |
| 220 PSHU - RV | ATGTCCATAAAGCACTCCTCACGC |
| 220 PSHU - FW | AGCAGCCTAACTTGGAGAATTCATATGGATGAT |
| 220 GBLOCK - RV | CAAGTTAGGCTGCTTCGCTACC |
| 224 GBLOCK - FW | AGGAGTGCTTTGAATCTATGCTTATTTTGTTGGGATAC |
| 224 PSHU - RV | TAAGCATAGATTCAAAGCACTCCTCACGC |
| 224 PSHU - FW | GAGGCAGCATAACTTGGAGAATTCATATGGATGATG |
| 224 GBLOCK - RV | CTCCAAGTTATGCTGCCTCCTCC |
| 259 GBLOCK - FW | GAGGAGTGCTTTATGGATTTGGCGACGC |
| 259 PSHU - RV | CCAAATCCATAAAGCACTCCTCACGC |
| 259 PSHU - FW | AGAAGGAGCATAACTTGGAGAATTCATATGGATGAT |
| 259 GBLOCK - RV | TTCTCCAAGTTATGCTCCTTCTGCTGC |
| 244 GBLOCK - FW | GGAGTGCTTTATGGACATGGCGACAATC |
| 244 PSHU - RV | CCATGTCCATAAAGCACTCCTCACGC |
| 244 PSHU - FW | AGAAGGAGCGTAACTTGGAGAATTCATATGGATGA |
| 244 GBLOCK - RV | TCCAAGTTACGCTCCTTCTGCGG |
| N41 GBLOCK - FW | GGAGTGCTTTATGCTTGTGATCATCGGG |
| N41 PSHU - RV | TCACAAGCATAAAGCACTCCTCACGC |

| | |
|---|---|
| N41 PSHU - FW | GGAAGAAGCATAACTTGGAGAATTCATATGGATGAT |
| N41 GBLOCK - RV | ATTCTCCAAGTTATGCTTCTTCCTCAGTGG |
| N65 GBLOCK - FW | AGGAGTGCTTTATGGATTTATCAACGATCATCGG |
| N65 PSHU - RV | GTTGATAAATCCATAAAGCACTCCTCACGC |
| N65 PSHU - FW | CTTGCGTGGATAACTTGGAGAATTCATATGGATGATG |
| N65 GBLOCK - RV | TCCAAGTTATCCACGCAAGGCC |
| 266 GBLOCK - FW | GGAGTGCTTTATCGCGACTATCATCGG |
| 266 PSHU - RV | TAGTCGCGATAAAGCACTCCTCACGC |
| 266 PSHU - FW | GCCGAGTGACTTGGAGAATTCATATGGATGATG |
| 266 GBLOCK - RV | TTCTCCAAGTCACTCGGCTTTCTCG |
| 332 GBLOCK - FW | AGGAGTGCTTTATTACAACGATTATTGGCTTAGTTTTAGG |
| 332 PSHU - RV | CAATAATCGTTGTAATAAAGCACTCCTCACGC |
| 332 PSHU - FW | AGCTGGAATGACTTGGAGAATTCATATGGATGATG |
| 332 GBLOCK - RV | ATTCTCCAAGTCATTCCAGCTTTTCGCG |
| 333 GBLOCK - FW | AGGAGTGCTTTATGTTTGCAATTATCGGGATCAT |
| 333 PSHU - RV | GATAATTGCAAACATAAAGCACTCCTCACGC |
| 333 PSHU - FW | CGTTCGAATGACTTGGAGAATTCATATGGATGATG |
| 333 GBLOCK - RV | AATTCTCCAAGTCATTCGAACGAGGGAC |
| 440 GBLOCK - FW | GGAGTGCTTTATCGCCACTATTATCGGATTG |
| 440 PSHU - RV | TAGTGGCGATAAAGCACTCCTCACGC |
| 440 PSHU - FW | AAGCAGAGTGACTTGGAGAATTCATATGGATGATGAA |
| 440 GBLOCK - RV | GAATTCTCCAAGTCACTCTGCTTTCTCGC |
| WT-VA178-79IG-F | CCCATTACAGCCCCAATAATACCAAACGCCGGAAGTG |
| WT-VA178-79IG-R | CACTTCCGGCGTTTGGTATTATTGGGGCTGTAATGGG |
| WT-I79G-F | CCCATTACAGCCCCAATAATACCAAACGCCGGAAGTG |

| | |
|---|---|
| WT-I79G-R | CACTTCCGGCGTTTGGTATTATTGGGGCTGTAATGGG |
| WT-Y217N-F | GGAAATAAATCCGTTAGCCAATAAAATGCCGAGGAAAGTC |
| WT-Y217N-R | GACTTTCCTCGGCATTTTATTGGCTAACGGATTTATTTCC |
| 180-VA178-79IG-F | AGGACTGCGCCAATAATGCCGAACGCGGGCA |
| 180-VA178-79IG-R | TGCCCGCGTTCGGCATTATTGGCGCAGTCCT |
| 180-Y217N-F | CAAGTAACCCGTTAGCCATGAAGACTCCCAGAAATG |
| 180-Y217N-R | CATTTCTGGGAGTCTTCATGGCTAACGGGTTACTTG |
| 220-VA178-79IG-F | CCCAAAACGGCACCTATAATTCCAAAGCCAGGCATACCAT |
| 220-VA178-79IG-R | ATGGTATGCCTGGCTTTGGAATTATAGGTGCCGTTTTGGG |
| 220-Y217N-F | GGACCAAAGAAGCCGTTCGCTGCCAAAATACCG |
| 220-Y217N-R | CGGTATTTTGGCAGCGAACGGCTTCTTTGGTCC |
| 333-VA178-79IG-F | CCCAATACCGCTCCTATAATCCCAAGAGCGGGTAAGCC |
| 333-VA178-79IG-R | GGCTTACCCGCTCTTGGGATTATAGGAGCGGTATTGGG |
| 333-Y217N-F | GGCCAACAAAGCCGTTGGCTAAGAACACGCC |
| 333-Y217N-R | GGCGTGTTCTTAGCCAACGGCTTTGTTGGCC |
| N41-VA178-79IG-F | CCCTAAAACGGCACCTATAATACCGAATCCGGGCAATCCG |
| N41-VA178-79IG-R | CGGATTGCCCGGATTCGGTATTATAGGTGCCGTTTTAGGG |
| N41-Y217N-F | AGGCCCAACAAATCCATTAGCAAGAAGGATTCCCA |
| NL 244 YG F | TGGCGAACGGAAGGAAAAGACCATAAGCGGAAAAGATGCCATAAAGGGTG |
| NL 244 YG R | CACCCTTTATGGCATCTTTTCCGCTTATGGTCTTTTCCTTCCGTTCGCCA |
| IG 244 VA F | GATCACCGCAGCAACAATACCCATGGTGGGGGC |
| IG 244 VA R | GCCCCCACCATGGGTATTGTTGCTGCGGTGATC |
| NM 259 YG F | AGCAATAGGAAAGAATACCCCATAACTCAGAATTGCACCATATAAGGTCGTAAGCAATGC |

| | |
|---|---|
| NM 259 YG R | GCATTGCTTACGACCTTATATGGTGCAATTCTGAGTTATGGGGTATTCTTTCCTATTGCT |
| IG 259 VA F | CCAAACCTACTAATGTTGCAACCATCCCCATGGCCGGT |
| IG 259 VA R | ACCGGCCATGGGGATGGTTGCAACATTAGTAGGTTTGG |
| NV 266 YG F | CGATGGGCAAAAAGATGCCATACGCACTTGCGACACCAT |
| NV 266 YG R | ATGGTGTCGCAAGTGCGTATGGCATCTTTTTGCCCATCG |
| IG 266 VA F | CGCGCCTACGATGGGTATTGTCGCAACTGTGATGGG |
| IG 266 VA R | CCCATCACAGTTGCGACAATACCCATCGTAGGCGCG |
| V 332 G F | CGAAGGGGCCAAATAAACCATACGCGGAGAAGATC |
| V 332 G R | GATCTTCTCCGCGTATGGTTTATTTGGCCCCTTCG |
| IG 332 VA F | GCCCCATTACAGCCGCAACGATCCCCATAGTCG |
| IG 332 VA R | CGACTATGGGGATCGTTGCGGCTGTAATGGGGC |
| NL 440 YG F | TGTTCGCGATGGGCAAAAAGATCCCATACGCAAGAATTGCCCCATACAATG |
| NL 440 YG R | CATTGTATGGGGCAATTCTTGCGTATGGGATCTTTTTGCCCATCGCGAACA |
| IG 440 VA F | CGCCCGCATTCGGTATGGTTGCAACACTTATTGGTCTTG |
| IG 440 VA R | CAAGACCAATAAGTGTTGCAACCATACCGAATGCGGGCG |
| 244-G40A-F | AACCGCTCCGATTGTAGCGCCGAAAACGATAAG |
| 244-G40A-R | CTTATCGTTTTCGGCGCTACAATCGGAGCGGTT |
| 244-F49N-F | CTTAATTTCATTCATTGGGTTAGAAATCAGAACCGCTCCGATTGTAC |
| 244-F49N-R | GTACAATCGGAGCGGTTCTGATTTCTAACCCAATGAATGAAATTAAG |
| 244-K262S,S263K-F | CTCATTGGGCGGAAGATACTTTCTCAATTTCTCCTCAATAACACGCG |

| | |
|---|---|
| 244-K262S,S263K-F | CGCGTGTTATTGAGGAGAAATTGAGAAAGTATCTTCCGCCCAATGAG |
| 259-G40A-F | TAAGGACAACAAAGATAGAAGCTCCCACTACGATAAGAATG |
| 259-G40A-R | CATTCTTATCGTAGTGGGAGCTTCTATCTTTGTTGTCCTTA |
| 259-F49N-F | GCTCCAAAAAATTGTCCCATAGTGTTCTTCATAAGGACAACAAAGATAGA |
| 259-F49N-R | TCTATCTTTGTTGTCCTTATGAAGAACACTATGGGACAATTTTTTGGAGC |
| 259-K262S,S263K-F | CGCGTTTACTCTCGTTTAAATACTTCCTCAAATAACTGTCAATCACGCG |
| 259-K262S,S263K-F | CGCGTGATTGACAGTTATTTGAGGAAGTATTTAAACGAGAGTAAACGCG |
| 266-G40A-F | TCGTGGCACCAATCGTGGCTCCAAATACAATGATG |
| 266-G40A-R | CATCATTGTATTTGGAGCCACGATTGGTGCCACGA |
| 266-F49N-F | TTCAGTTCTTTCAGAGGGTTCGATACCATCGTGGCACC |
| 266-F49N-R | GGTGCCACGATGGTATCGAACCCTCTGAAAGAACTGAA |
| 266-K262S,S263K-F | GCGCTCCTTTGGTGGTAAAAACTTTCTTAATTTTTCTTCGATGATGCGTGGATTTTCACC |
| 266-K262S,S263K-F | GGTGAAAATCCACGCATCATCGAAGAAAAATTAAGAAAGTTTTTACCACCAAAGGAGCGC |
| 332-G40A-F | TCACAGCACCGATAGTCGCGCCAAAAATGATAATG |
| 332-G40A-R | CATTATCATTTTTGGCGCGACTATCGGTGCTGTGA |
| 332-F49N-F | TTCCTTCATGGGATTTGACACCATCACAGCACCGATAGTC |
| 332-F49N-R | GACTATCGGTGCTGTGATGGTGTCAAATCCCATGAAGGAA |
| 332-K262S,S263K-F | TTCTTTGGGGCTTAAGTACTTCCTTAACTTTTCTTCGATTACACGCGGGTTTTCGCC |
| 332-K262S,S263K-F | GGCGAAAACCCGCGTGTAATCGAAGAAAAGTTAAGGAAGTACTTAAGCCCCAAAGAA |

| | |
|---|---|
| 266-K262S,S263K-F | GCGCTCCTTTGGTGGTAAAAACTTTCTTAATTTTTCTTCGATGATGCGTGGATTTTCACC |
| 266-K262S,S263K-F | GGTGAAAATCCACGCATCATCGAAGAAAAATTAAGAAAGTTTTTACCACCAAAGGAGCGC |
| 332-G40A-F | TCACAGCACCGATAGTCGCGCCAAAAATGATAATG |
| 332-G40A-R | CATTATCATTTTTGGCGCGACTATCGGTGCTGTGA |
| 332-F49N-F | TTCCTTCATGGGATTTGACACCATCACAGCACCGATAGTC |
| 332-F49N-R | GACTATCGGTGCTGTGATGGTGTCAAATCCCATGAAGGAA |
| 332-K262S,S263K-F | TTCTTTGGGGCTTAAGTACTTCCTTAACTTTTCTTCGATTACACGCGGGTTTTCGCC |
| 332-K262S,S263K-F | GGCGAAAACCCGCGTGTAATCGAAGAAAAGTTAAGGAAGTACTTAAGCCCCAAAGAA |
| Plug Del Fs | CTAATAGTGAAAGCCCAATTCCCGG |
| Plug Del F-T | ACGGGCGGCGATCGCATTTCTAATAGTGAAAGCCCAATTCCCGG |
| Plug Del Rs | GGAGATGGAGATCAGCCACATCA |
| Plug Del R-T | AAATGCGATCGCCGCCCGTGGAGATGGAGATCAGCCACATCA |

**PCR conditions for linear vector preparation:**

| Step | Temperature | Time |
|---|---|---|
| Initial Denaturation | 98°C | 2 mins |
| X 25 Cycles | 98°C<br>65°C<br>72°C | 30 Sec<br>1 min<br>3 mins 30 Sec |
| | 72°C | 5 mins |

**PCR conditions for MotA insert preparation:**

| Step | Temperature | Time |
|---|---|---|
| Initial Denaturation | 98°C | 2 mins |
| X 15 Cycles | 98°C<br>65°C<br>72°C | 30 Sec<br>1 min<br>3 mins 30 Sec |
| | 72°C | 5 mins |
